# Natural antibodies as “eat-me” signals for phagocytosis of necrotic cell debris at sites of tissue injury

**DOI:** 10.1101/2023.03.29.533912

**Authors:** Matheus Silvério Mattos, Sofie Vandendriessche, Sara Schuermans, Lars Feyaerts, Nadine Hövelmeyer, Ari Waisman, Pedro Elias Marques

## Abstract

Natural antibodies (NAbs) are circulating polyreactive immunoglobulins that bind endogenous and exogenous antigens. Here, we investigated the role of NAbs in driving the clearance of necrotic cell debris from injury sites. Using mouse models of liver injury, we observed that IgM and IgG NAbs opsonize necrotic debris *in vivo* by recognizing common self-molecules such as histones, actin, phosphoinositides and cardiolipin, but not phosphatidylserine. Importantly, mice lacking NAbs presented impaired recovery from liver injury, which was correlated to sustained presence of necrotic debris in the tissue, prolonged inflammation and reduced hepatocellular proliferation. Mechanistically, necrotic debris phagocytosis was dependent on NAbs *in vitro* and *in vivo*, and restitution with total immunoglobulins rescued the defective recovery from liver injury in immunodeficient mice. In summary, we showed that NAbs opsonize necrotic cell debris and act as “eat-me” signals for engulfment through FcγRs and CD11b, driving the recovery from tissue injury.

**Highlights:**

- Natural antibodies opsonize exposed self-antigens upon necrotic cell death.
- The phagocytosis of necrotic cell debris requires natural antibodies, FcγRs and CD11b.
- Natural antibodies drive cellular proliferation and tissue regeneration after liver injury.
- Treatment with natural antibodies improves the recovery from liver injury in both immunodeficient and immunocompetent mice.

**Graphical abstract:** 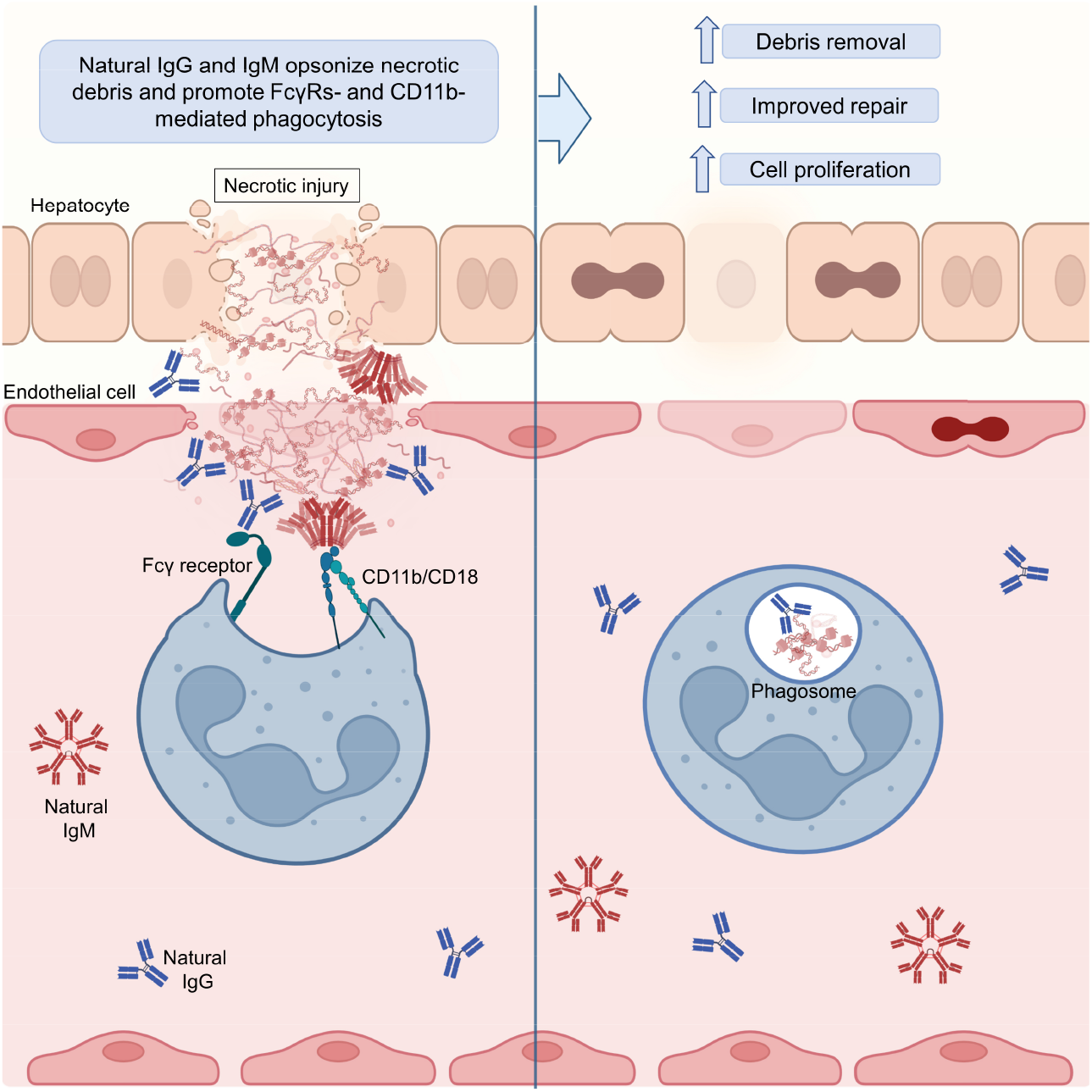

## Introduction

Natural antibodies (NAbs) are circulating polyreactive immunoglobulins that arise without exogenous antigenic stimulation. NAbs are mostly IgM and IgG isotypes produced mainly by CD5^+^ B-1 cells^1^. NAbs are characterized as low affinity antibodies that, due to their polyreactivity, bind to different classes of self and exogenous molecules including proteins, nucleic acids, carbohydrates, lipids or combinations thereof^2^. In contrast to adaptive antibodies that can bind virtually to any epitope, NAbs have germline-encoded variable regions which restrict their recognition capacity to phylogenetically conserved molecules^3,4^. In fact, natural IgM and IgG harvested from human cord blood or germ-free mice can bind to common molecules such as DNA, fibrin, actin, tubulin, lysophosphatidylcholine and oxidized phospholipids^5–8^. Since the term “natural antibodies” was first introduced in the early 1960s^9^, NAbs were neglected because the existence of self-binding antibodies is contradictory with Burnet’s clonal selection theory^10^, causing the biological function of NAbs to remain largely elusive. Indeed, although some autoimmune diseases are associated with autoantibodies that breach Burnet’s clonal selection theory, it is equally evident that not all antibodies that react to self-molecules are pathogenic, since circulating NAbs are constitutively present from birth to adulthood. Thus, it is reasonable to postulate that NAbs have a physiological role throughout life.

Necrotic cell death is inexorably connected to human life as a result of our daily behavior, since we are constantly exposed to stresses such as burns, traumas and intoxications that culminate in necrotic injuries. Necrotic cell death, accidental or programmed, and regardless of the wide spectrum of necrosis-initiating events, converges in plasma membrane rupture and the consequent release of cellular content (debris) in the tissue^11^. Once exposed, necrotic debris are recognized as damage-associated molecular patterns (DAMPs), acting as powerful inducers of inflammation. Not surprisingly, the generation and longevity of necrotic debris in tissues are associated with chronic inflammation and worsening of atherosclerosis, arthritis, liver injury, systemic lupus erythematosus and neurodegenerative disorders^12,13^. In order to avoid this grim prospect, necrotic debris must be efficiently cleared from tissues.

The clearance of dead cells is mainly performed by professional phagocytes, in which efferocytosis, the phagocytosis of apoptotic cells, has been extensively described^14^. During apoptosis, intracellular components are packed into sealed apoptotic bodies expressing phosphatidylserine (PS), a central “eat-me” signal^15^ recognized by phagocytic receptors such as MerTK, BAI-1, TIM-(1-4) and CD36, among others^16–19^. The mechanisms involved in the recognition and phagocytosis of necrotic cells, however, are largely unexplored. Given to its broad molecular and size heterogeneity, the recognition and phagocytosis of necrotic debris represents a major challenge to the limited range of leukocyte phagocytic receptors. A way to bypass the need of multiple receptors would be to use polyreactive adaptor molecules bridging the recognition of necrotic debris by phagocytes. In this context, polyreactive NAbs may enable the binding and internalization of necrotic debris through Fc receptors (FcRs), which recognize the Fc-portion of immunoglobulins in antibody-coated targets. FcγRs and Fc-α/μ were shown to mediate phagocytosis of diverse IgG- and IgM-opsonized targets, respectively^20,21^. Importantly, NAbs were shown to bind to necrotic cells *in vitro* and *in vivo*^5,22^. IgM and IgG binding may also lead to the activation of the complement cascade, which was implicated in the clearance of late apoptotic and necrotic cells *in vitro*^13,23,24^.

Here, we utilized *in vitro* and *in vivo* approaches to understand how necrotic cells are cleared from injured tissue. Protein and lipid blots were performed using purified debris to determine the (multiple) cellular components in necrotic debris that were recognized by NAbs. We used liver intravital microscopy to image the phagocytosis of necrotic cells debris in real time, assessing the participation of natural antibodies in both immunocompetent and immunodeficient (RAG2^-/-^ and IgMi) mice. We further analyzed the phagocytosis of NAbs-opsonized necrotic debris *in vitro* in primary human and murine neutrophils in order to identify receptors involved in the NAbs-mediated cell debris clearance. Our study showed that NAbs are specific adaptors for the recognition and removal of necrotic cell debris from injury sites.

## Results

### Natural IgM and IgG are rapidly deposited on necrotic cell debris following injury

To evaluate the role of NAbs in necrotic cell debris clearance, we used a well-described and clinically relevant model of acute liver necrosis, namely drug-induced liver injury by acetaminophen (APAP) overdose^25,26^. Consistent with previous work, mice that received 600 mg/kg of APAP orally developed extensive and time-dependent hepatocellular necrosis starting 6h post-overdose, peaking at 12h-24h and resolving from 48h onwards, as assessed by fibrin(ogen) deposition in liver cryosections (**Figure 1A, 1B and supplementary figure 1**). Fibrin(ogen) staining was used as the main method for identification and measurement of liver necrotic areas. Liver injury was confirmed by increased levels of alanine transaminase (ALT) in murine sera after the APAP challenge (**Figure 1C**), which followed similar kinetics to the centrilobular necrosis observed in cryosections. Liver injury resulted in recruitment of neutrophils to the liver as early as 6h (**Figure 1D**), followed by significant recruitment of monocytes 12h after APAP intoxication (**Figure 1E**). The recruitment of both types of phagocytes peaked at 24h, however, the percentage of monocytes remained elevated until 72h, further into the resolution phase of injury (**Figure 1D and 1E**).

**Figure 1:**
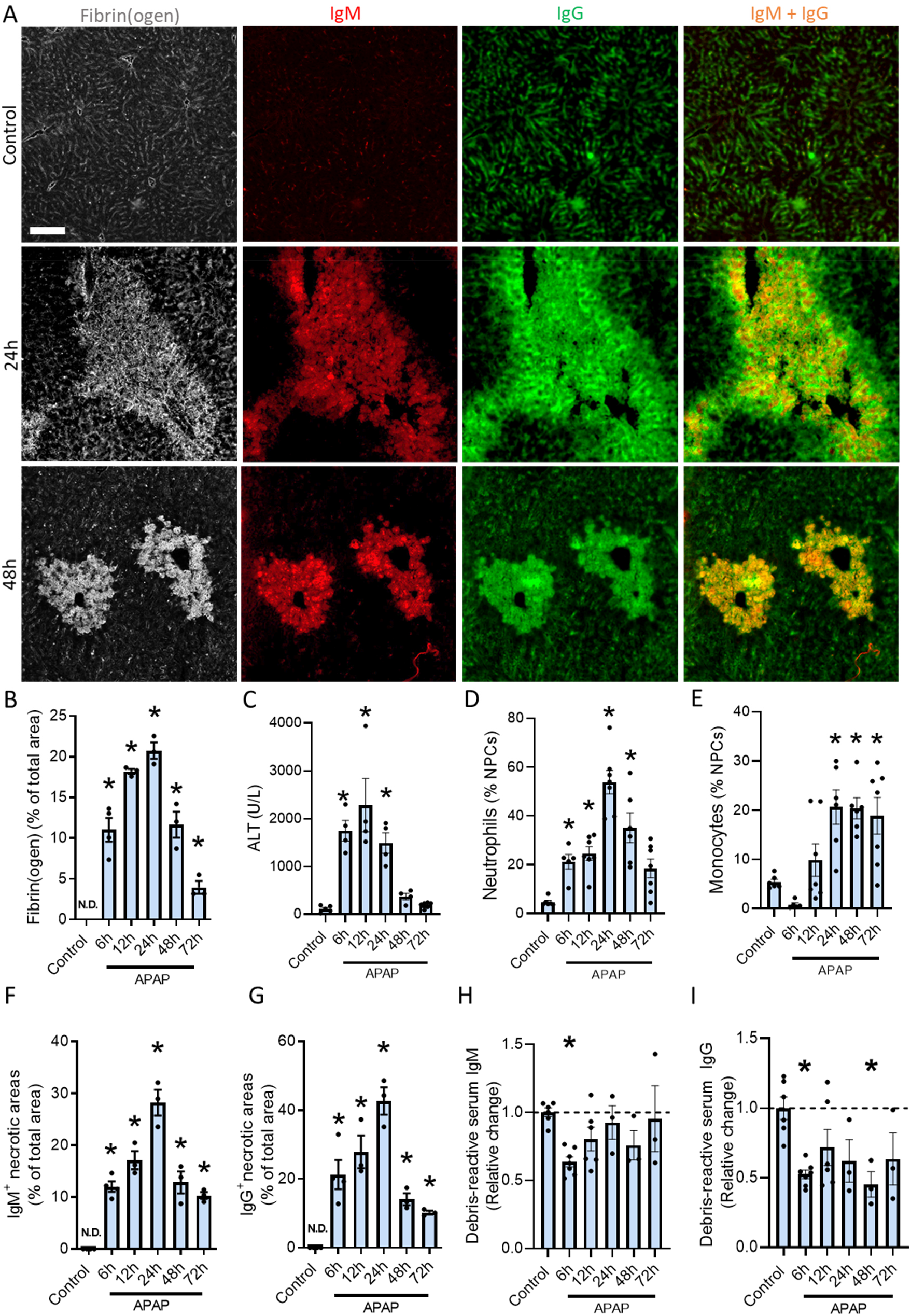
Natural IgM and IgG are rapidly deposited on necrotic cells following injury. (**A**) Representative images showing immunostaining of liver cryosections from control mice and mice gavaged with APAP (600 mg/kg) for 24 and 48 hours. Gray: fibrin(ogen), red: IgM, green: IgG, orange: merged IgM and IgG. Scale bar = 100 μm. See full panel from 6 to 72 hours in Figure S1. (**B**) Quantification of the fibrin(ogen) positive area fraction in liver cryosections. (**C**) Serum ALT in APAP challenged mice. (**D**) Flow cytometry of liver non parenchymal cells (NPCs) showing the percentage of neutrophils (Ly6G^+^). (**E**) Flow cytometry of liver NPCs showing the percentage of monocytes (Ly6G^-^/CCR2^+^). (**F**) Quantification of the IgM positive area fraction in liver cryosections. (**G**) Quantification of the IgG positive area fraction in liver cryosections. (**H**) Quantification of debris-reactive serum IgM after APAP challenge. (**I**) Quantification of debris-reactive serum IgG after APAP challenge. Data are from mice 6-72 hours after an oral gavage of 600 mg/kg APAP (**A-I**) and are represented as mean ± SEM. Each dot in the graph represents a single mouse (**B-I**). Image quantifications were pooled from 10 fields of view (**B, F, G**). * p< 0.05 compared to control group.

Once the kinetics of hepatic necrosis was ascertained, we investigated if NAbs were deposited in the injured liver. Liver cryosections were stained with fluorescently labeled anti-mouse IgM and anti-mouse IgG antibodies, which showed that both natural IgM and IgG were bound to the necrotic areas identified by the fibrin(ogen) staining (**Figure 1A and supplementary figure 1**). Interestingly, accumulation of IgM and IgG in necrotic areas occurred in parallel to the development of liver injury, also peaking at 24h and decreasing with tissue recovery 48-72h after injury (**Figures 1A, 1F, 1G and supplementary figure 1**). We also found that the deposition of NAbs in the necrotic liver led to the reduction of circulating IgM and IgG that is reactive to necrotic debris (**Figure 1H and 1I**). While both NAbs isotypes were reduced in the bloodstream 6h after APAP, the reduction of natural IgG was more pronounced and persistent during liver injury. This shows that these antibodies were sequestered from the bloodstream as they became deposited in injury sites.

To visualize the binding of NAbs to necrotic cells in living tissue, confocal intravital microscopy (IVM) was performed 24h after the APAP challenge. Prior to imaging, mice were injected intravenously (i.v.) with Sytox green, a membrane-impermeable DNA dye, and anti-mouse IgM and IgG to label the necrotic cells and deposited NAbs, respectively. Using this approach, we confirmed the binding of both natural IgM and IgG to necrotic cells *in vivo* (**Figure 2A and supplementary video 1**). Moreover, plasma membrane of dying hepatocytes were often positive for IgM/IgG labeling even before these cells becoming positive for Sytox green, indicating that the binding of NAbs is fast and independent of the full demise of the cells (**Supplementary video 2**). Altogether, these data demonstrate that during liver injury NAbs leave the systemic circulation and become deposited within the necrotic areas. Binding of NAbs to necrotic debris followed the kinetics of liver injury, decreasing progressively towards 72h after the APAP challenge. These observations suggested that recognition and binding of NAbs to necrotic debris may have a role in necrotic cell clearance and tissue repair.

**Figure 2:**
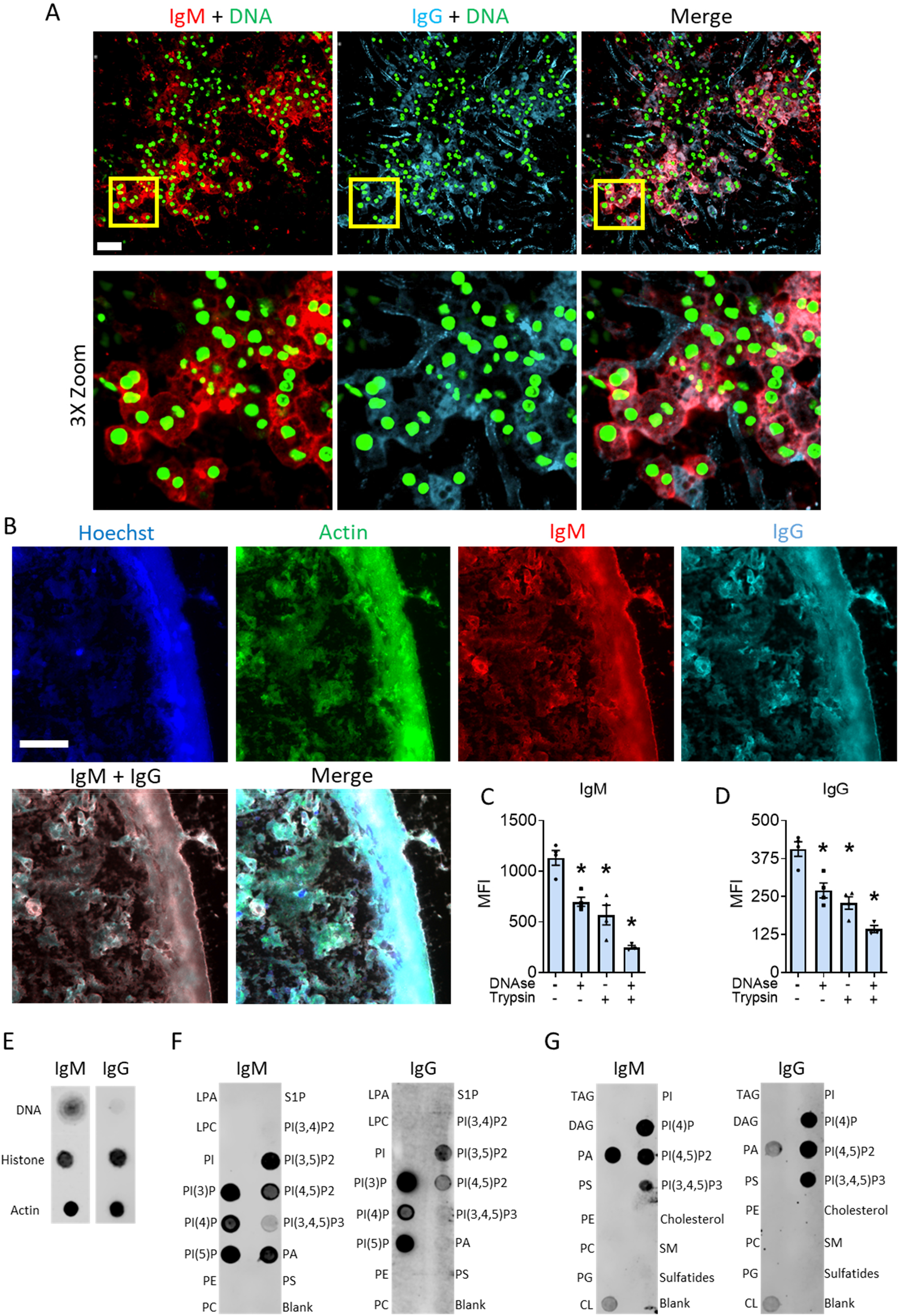
Natural IgM and IgG antibodies bind multiple self-antigens exposed upon necrotic cell death. (**A**) Representative intravital microscopy images showing IgM (red) and IgG (cyan) deposition in dead/dying hepatocytes (sytox green^+^) 24h after APAP intoxication (600 mg/kg). Scale bar = 50 μm. See also supplementary videos 1 and 2. (**B**) Binding of serum natural IgM (red) and IgG (cyan) to a necrotic hepatocyte debris spot. DNA is in blue and f-actin in green. Scale bar = 100 μm. (**C and D**) Mean fluorescence intensity (MFI) of IgM and IgG labeling after pre-treatment of debris spot for 15 minutes with DNase and/or Trypsin before adding serum. Data are represented as mean ± SEM. *p< 0.05 compared to untreated samples. (**E**) Dot blots showing the reactivity of purified IgM and IgG to 4 μg of purified DNA, histones and actin. (**F and G**) Lipid blot showing the reactivity of purified IgM and IgG to lipids (100 pmol). CL: cardiolipin; DAG: diacylglycerol; LPA: lysophosphatidic acid; LPC: lysophosphatidylcholine; PA: phosphatidic acid; PC: phosphatidylcholine; PE: phosphatidylethanolamine; PG: phosphatidylglycerol; PI; phosphatidylinositol; PI(3)P: PI 3-phosphate; PI(4)P: PI 4-phosphate; PI(5)P: PI 5-phosphate; PI(3,4)P2: PI 3,4-bisphosphate; PI(4,5)P2: PI 4,5-bisphosphate; PI(3,4,5)P3: PI 3,4,5-trisphosphate; PS: phosphatidylserine; S1P: sphingosine 1-phosphate; SM: sphingomyelin; TAG: triacylglycerol Data are represented as mean ± SEM. *p< 0.05 compared to the untreated sample.

### Natural IgM and IgG antibodies bind multiple self-antigens exposed upon necrotic cell death

We next sought to investigate if NAbs bound specifically to necrotic debris or if they were trapped nonspecifically in injury sites. For this purpose, hepatocytes from healthy mouse livers were purified and crushed mechanically to expose their intracellular contents, which were then spotted onto coverslips. The necrotic debris spots were incubated with mouse serum and labeled with anti-mouse IgM and IgG. We observed that the hepatocyte debris spots, shown by DNA and F-actin staining, were bound by IgM and IgG from healthy mouse serum (**Figure 2B**). The binding of antibodies was antigen-specific, since incubation with secondary antibodies alone yielded no labelling of the debris (**Supplementary figure 2A**). Next, the types of antigens that these polyreactive NAbs recognized in the necrotic debris were investigated. To distinguish whether NAbs were preferentially bound to nucleic acids or proteins, we pre-treated the debris spots with DNase and/or trypsin for 15 minutes. Treatment with DNase and trypsin was sufficient to degrade DNA and F-actin substantially, but not completely (**Supplementary figure 2B**). Pre-incubation of debris spots with either of the enzymes reduced both IgM and IgG labelling significantly, suggesting that NAbs bind DNA and protein epitopes (**Figure 2C and 2D**). Combined treatment with DNAse and trypsin resulted in a cumulative reduction of IgM and IgG binding. To further investigate the specificity of IgM and IgG NAbs towards necrotic cell debris, dot blots with purified DNA, histones and actin were performed and incubated with mouse serum. We found that serum NAbs indeed bind directly to histones, actin and DNA, the latter albeit with lower avidity (**Supplementary figure 2C**). In order to avoid interference from other possible debris-binding molecules present in serum, we also performed dot blots using purified murine IgM or IgG and we found a similar binding pattern (**Figure 2E**).

To assess if the recognition of necrotic cell molecules by NAbs also encompassed lipids, we performed assays with lipid strips containing a variety of cellular phospholipids, phosphoinositides and their intermediates. We observed that IgM and IgG NAbs recognized several phosphoinositides such as phosphatidylinositol 3 phosphate [PI(3)P], PI(4)P, PI(5)P, PI(3,5)P2, PI(4,5)P2 (**Figure 2F**) as well as phosphatidic acid (PA), PI(3,4,5)P3 and the mitochondrial phospholipid cardiolipin (CL) (**Figure 2G**). Importantly, we observed no binding of IgM nor IgG NAbs to phosphatidylserine (PS), phosphatidylethanolamine (PE), phosphatidylcholine (PC), phosphatidylglycerol (PG), phosphatidylinositol (PI), sphingomyelin (SM), sphingosine-1-phosphate (S1P), diacylglycerol (DAG), lysophosphatidic acid (LPA), lysophosphocholine (LPC), the triglyceride glyceryl tripalmitate (TAG) or cholesterol. These data indicate that NAbs are capable of recognizing specific categories of membrane lipids, e.g. phosphoinositides and cardiolipin, which are restricted to the cytoplasmic leaflet of the plasma membrane and to intracellular organelles such as endosomes, Golgi, endoplasmic reticulum and mitochondria (for cardiolipin)^27,28^. These results further show that both IgM and IgG NAbs bind to multiple classes of cellular components that are exposed after necrotic cell death including DNA, actin, histones and phospholipids. Yet, IgM and IgG are able to bind essentially to the same antigens, suggesting a redundant role for the two types of NAbs.

### Mice that lack NAbs have impaired cell debris clearance, prolonged inflammation and delayed recovery from liver injury

Once we determined that NAbs recognize necrotic cell debris and are present within the necrotic areas *in vivo*, we investigated the hypothesis that NAbs play a role in debris clearance and recovery from injury. For this, we challenged RAG2^-/-^ mice, which lack mature lymphocytes and consequently antibodies, with APAP for 24h and 48h. We chose to work with these 2 timepoints to evaluate if there were differences in the peak of injury (24h) and to assess if a defective tissue repair would occur (48h). Immunostaining of liver cryosections of APAP-challenged mice were performed to confirm the absence of NAbs deposition in the injured livers of RAG2^-/-^ mice. As expected, there was no IgM nor IgG present in RAG2^-/-^ livers 24h or 48h after APAP intoxication, in contrast to wild-type (WT) RAG2^+/+^ samples (**Supplementary Figure 3**).

To evaluate the hepatic necrotic areas directly, we performed IVM of livers of WT and RAG2^-/-^ mice after APAP intoxication. The necrotic debris was identified with Sytox green, since DNA is a common debris component found in the liver following hepatocellular necrosis^11^. With this approach, we found that the DNA debris in livers of WT mice had already been largely cleared 48h after APAP, however, RAG2^-/-^ livers still presented a significant amount of extracellular DNA in the tissue, indicating an impairment in DNA debris clearance (**Figure 3A and 3B**). Fibrin(ogen) staining showed very similar results, in which RAG2^-/-^ mice presented significantly larger necrotic areas 48h post injury (**Figure 3C**).

**Figure 3:**
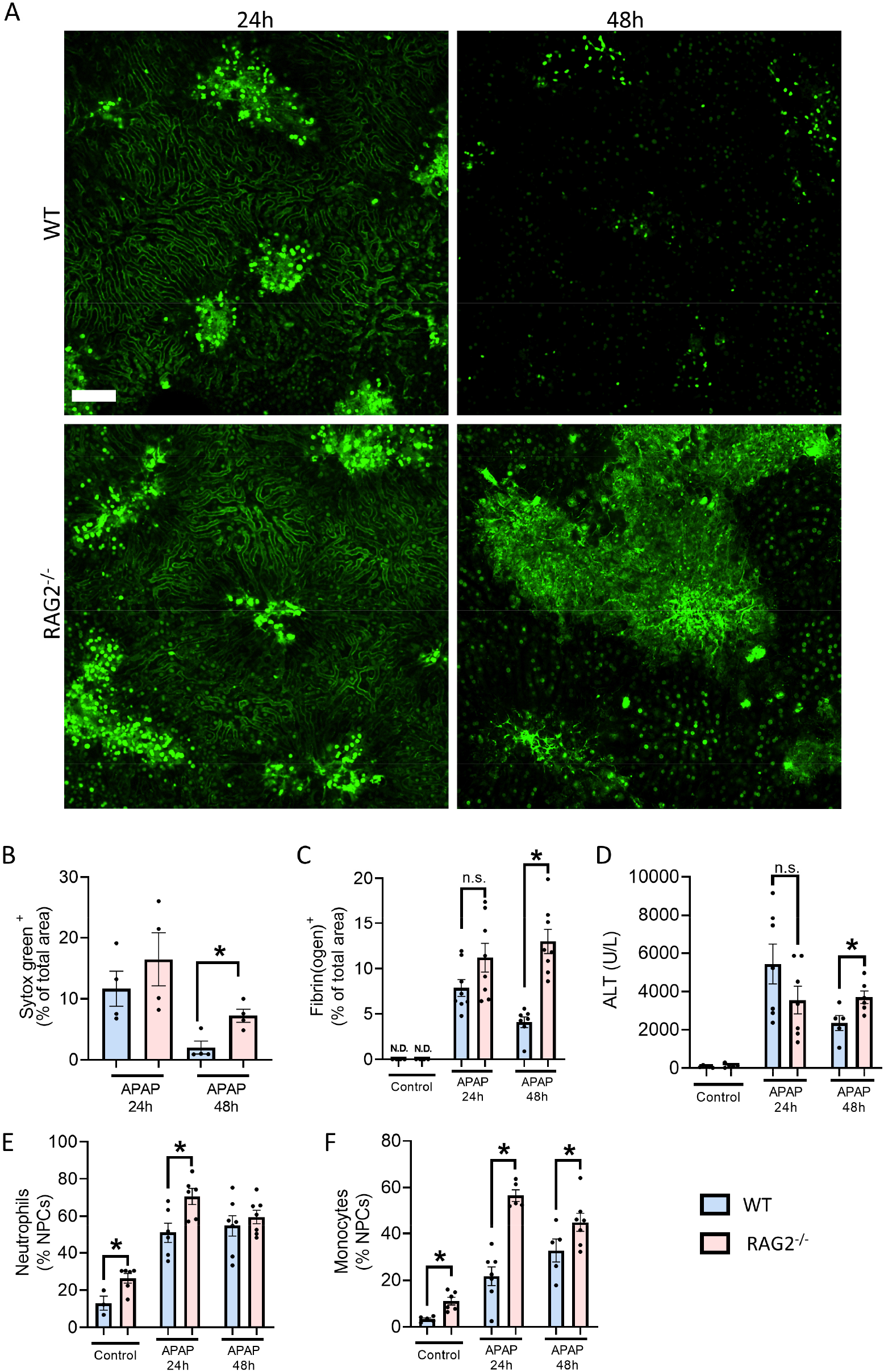
Mice that lack NAbs have impaired cell debris clearance, prolonged inflammation and delayed recovery from liver injury. (**A**) Representative intravital microscopy images of WT and RAG2^-/-^ mice showing the deposition of DNA (sytox green^+^) in the liver 24 and 48 hours after APAP overdose (600 mg/kg). Scale bar = 100 μm. (**B**) Quantification of the sytox green^+^ area in the liver. (**C**) Quantification of the fibrin(ogen)^+^ area fraction in liver cryosections. See supplementary figure 3A for IgM and IgG stainings. (**D**) Serum ALT levels in WT and RAG2^-/-^ mice 24 and 48 hours after APAP overdose. (**E**) Flow cytometry of liver non parenchymal cells (NPCs) showing the percentage of neutrophils (Ly6G^+^). (**F**) Flow cytometry of liver NPCs showing the percentage of monocytes (Ly6G^-^/CCR2^+^). Data are from WT and RAG2^-/-^ mice 24 and 48 hours after an oral gavage of 600 mg/kg APAP (**A-F**) and are represented as mean ± SEM. Each dot in the graph represents a single mouse (**B-F**). Image quantifications were pooled from 10 fields of view (**B-C**). *p< 0.05.

It is important to mention that the absence of NAbs did not cause excessive injury, as shown by similar ALT levels between WT and RAG2^-/-^ mice 24h after APAP (**Figure 3D**). Instead, RAG2^-/-^ mice presented ALT levels that remained as high in 48h as they were at the peak of injury, showing that liver injury is essentially not resolving. Fibrin(ogen) staining of RAG2^-/-^ livers reproduced the observations with DNA debris by IVM, in which trends to accumulated necrotic debris and larger necrotic areas 24h after APAP became significantly increased 48h after the challenge (**Figures 3C, 3D and supplementary figure 3A**). Altogether, RAG2^-/-^ mice presented a severe delay in liver recovery 48h post injury, as shown by DNA debris accumulation *in vivo*, larger fibrin(ogen)^+^ necrotic areas still present in the liver, and persistently elevated ALT levels (**Figures 3B-D**). Corroborating the prolonged liver injury of RAG2^-/-^ mice, we also observed a significantly higher recruitment of neutrophils and monocytes, indicating that RAG2^-/-^ livers are proportionally more inflamed when compared to WT (**Figures 3E and 3F**). Altogether, this suggests that RAG2^-/-^ mice have impaired necrotic debris clearance and a delayed resolution of liver injury.

### Mice that lack soluble antibodies (IgMi) have a defective regenerative response to necrotic liver injury

In order to refine our observations with RAG2^-/-^ mice, we investigated APAP-induced liver injury in IgMi mice. These mice develop mature lymphocytes, including B cells that express membrane IgM as the B cell receptor, however, they are unable to perform class switch and to differentiate into plasma cells^29^. The resultant phenotype is that IgMi mice have mature lymphocytes but do not secrete soluble antibodies^30^. To confirm the absence of NAbs in IgMi mice, we performed liver cryosections 24h and 48h after APAP overdose and stained it with fluorescent anti-IgM and anti-IgG antibodies. As in RAG2^-/-^ mice, we did not detect any IgM or IgG bound to the necrotic areas (**Figure 4A**). IgMi mice presented larger fibrin(ogen) deposition in the liver 24h after APAP, which remained larger at 48h when compared to WT mice (**Figure 4A and 4B**). Moreover, serum ALT levels were not different between IgMi and WT mice 24h after injury, however, ALT levels in IgMi mice remained significantly elevated at the 48h timepoint (**Figure 4C**), indicating that the necrotic injury is prolonged comparably in IgMi mice and RAG2^-/-^ mice.

**Figure 4:**
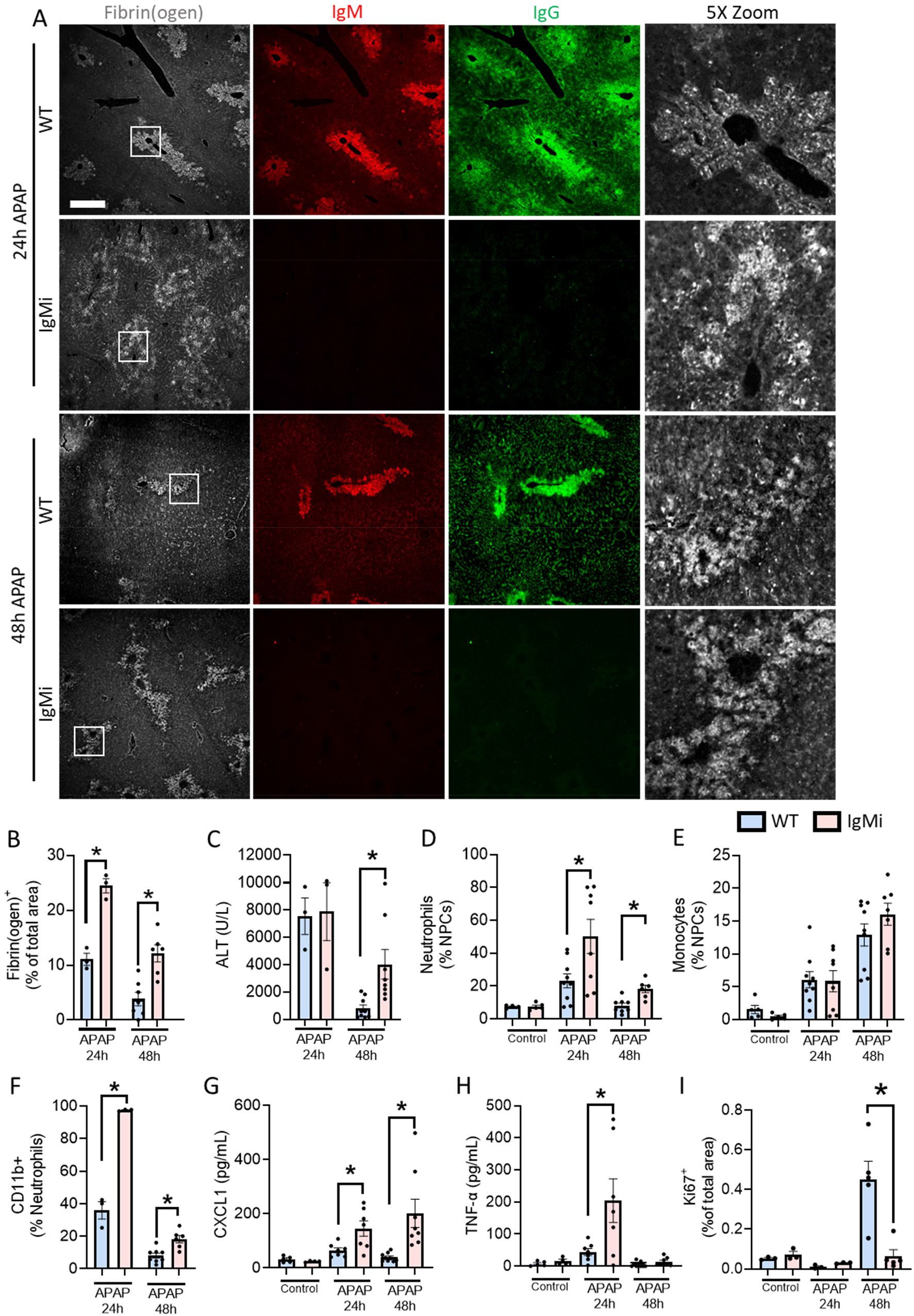
Mice that lack soluble antibodies (IgMi) have a defective regenerative response to necrotic liver injury. (**A**) Representative images showing immunostaining of liver cryosections from WT and IgMi mice, 24 and 48 hours after oral gavage with APAP (600 mg/kg). White squares indicate the area zoomed. Gray: fibrin(ogen), red: IgM and green: IgG. Scale bar = 100 μm. (**B**) Quantification of fibrin(ogen)^+^ area fraction in liver cryosections. (**C**) Serum ALT levels in WT and IgMi mice challenged with APAP. (**D**) Flow cytometry of liver non parenchymal cells (NPCs) showing the percentage of neutrophils (Ly6G^+^). (**E**) Flow cytometry of liver NPCs showing the percentage of monocytes (Ly6G^-^/CCR2^+^). (**F**) Percentage of neutrophils expressing CD11b by flow cytometry. (**G**) Serum levels of CXCL1 in control mice or after receiving APAP for 24 and 48 hours. (**H**) Serum levels of TNF-α as in G. See also figure S4 for IL-6 and IL-10 levels. (**I**) Quantification of Ki67^+^hepatocytes in liver cryosections in of control and APAP-challenged mice. Data are from WT and IgMi mice 24 and 48 hours after an oral gavage with 600 mg/kg APAP (A-I) and are represented as mean ± SEM. Each dot in the graph represents a single mouse (**B-I**). Image quantifications were pooled from 10 fields of view (**B, I**). *p< 0.05.

We also analyzed the recruitment of phagocytes to the livers of IgMi mice using flow cytometry. The migration of neutrophils to IgMi livers was significantly higher at both 24h and 48h post injury in comparison to WT mice (**Figure 4D**). Monocytes were also recruited to the livers of WT and IgMi mice throughout the injury, but no differences were observed between the mouse strains (**Figure 4E**). Interestingly, almost 100% of neutrophils in the livers of IgMi mice expressed surface CD11b 24h after injury, compared to only 40% in WT mice. CD11b expression is used as a marker of neutrophil activation, which indicated that hepatic neutrophils were more activated in IgMi mice during liver injury. Although CD11b expression decreased at 48h in both groups, it remained significantly higher in IgMi neutrophils (**Figure 4F**). In addition, we found increased levels of circulating CXCL1, TNF-α, IL-6 and IL-10 in IgMi mice, confirming that the absence of NAbs during necrotic liver damage promotes a severe dysregulation of the immune response to injury (**Figures 4G, 4H and Supplementary 4A, 4B**).

Of note, CXCL1 and TNF-α were significantly elevated during the peak of liver injury and CXCL1 remained elevated up to 48h in IgMi mice, which correlates to the disproportionate neutrophil recruitment and demonstrates the excessive pro-inflammatory state of mice lacking NAbs during injury.

We also hypothesized that besides restricting the extent of hepatic damage and inflammation during injury, NAbs had a beneficial effect on liver regeneration^22^. To evaluate this, we assessed the levels of cellular proliferation in the injured liver via immunostaining of Ki67 in cryosections of WT and IgMi mice. We found a significant increase in cellular proliferation 48h post injury (regenerative phase) in WT mice, which was essentially absent in the livers of NAbs-deficient IgMi mice (**Figure 4I**). This showed that NAbs drive hepatocellular proliferation after injury, which is required for liver regeneration and recovery. Altogether, the data on IgMi mice corroborated our findings in RAG2^-/-^ mice and confirmed that the defective liver recovery is mainly due to the absence of NAbs. The lack of NAbs worsens several critical events during liver injury, such as delaying the clearance of necrotic debris and the resolution of inflammation, but also by impairing the proliferation of hepatocytes.

### Restitution of NAbs to RAG2^-/-^ mice improves the recovery from liver injury

To confirm the role of NAbs in the recovery from liver injury, we subjected RAG2^-/-^ mice to APAP-induced liver injury and after 4h they received an i.v. transfer of RAG2^+/+^ (WT) or RAG2^-/-^ serum (lacking NAbs). Treatment with WT but not with RAG2^-/-^ serum successfully restored the deposition of natural IgM and IgG in necrotic areas in the liver (**Figure 5A**). The restitution of NAbs did not affect the initial injury significantly, as evidenced by similar fibrin(ogen)^+^ areas and serum ALT levels after 24h (**Figures 5A–5C**). It is important to mention that the serum ALT levels and fibrin(ogen) areas followed a similar pattern to what we observed in WT with RAG2^-/-^ mice (**Figure 3C and 3D**). The serum transfer did not affect the recruitment of neutrophils nor monocytes to the liver (**Supplementary figure 5A and 5B**), nor the production of CXCL1, TNF-α, IL-6 and IL-10, except for an increase in IL-10 48h after injury in mice treated with RAG2^-/-^ serum (**Supplementary figure 5C-F**). This indicates that the induction of injury and inflammation in the liver was not altered significantly by the serum transfer and that NAbs do not regulate leukocyte recruitment to the necrotic liver.

**Figure 5:**
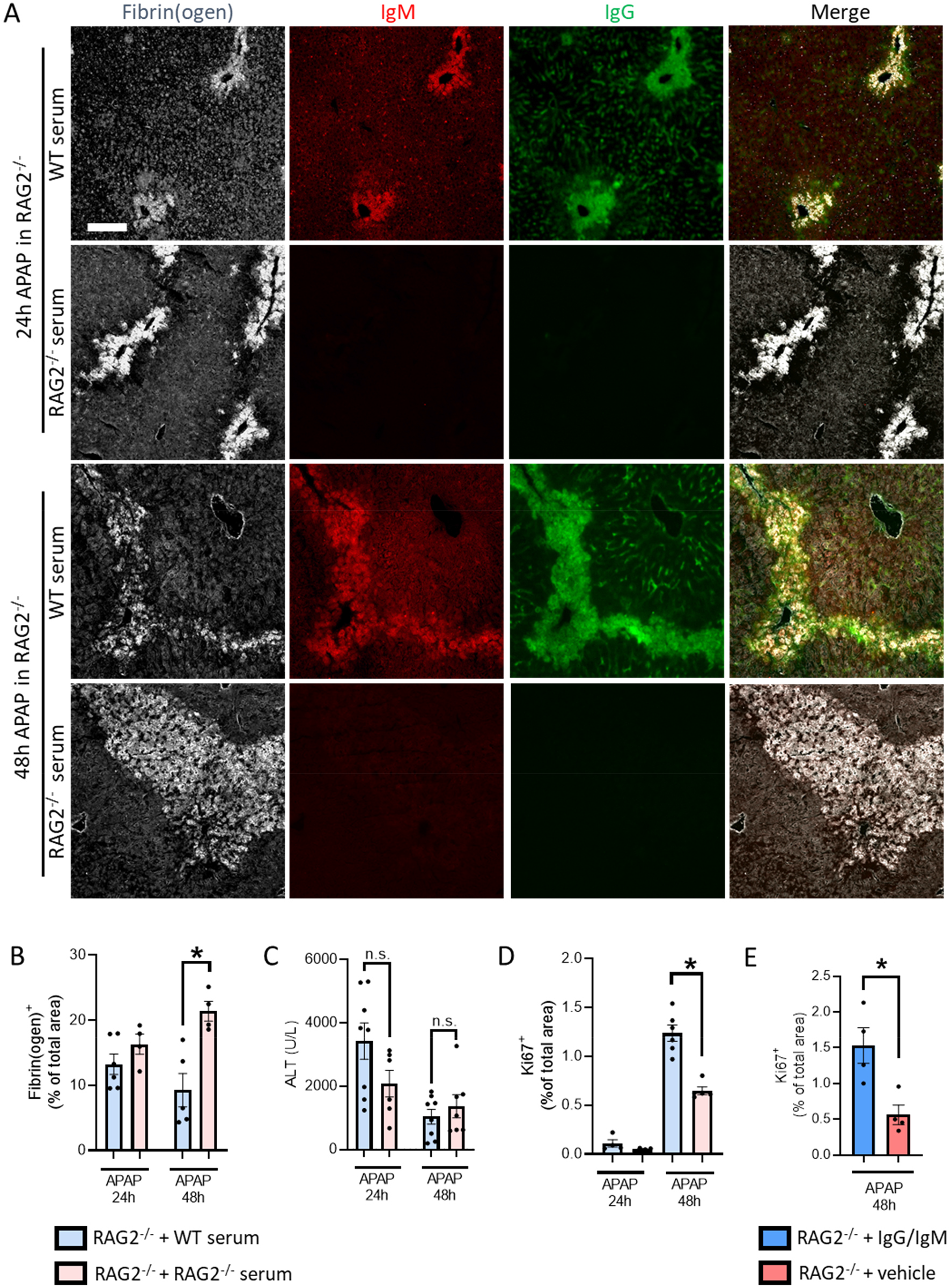
Restitution of NAbs to RAG2^-/-^ mice improves the recovery from liver injury. (**A**) Representative images showing immunostaining of liver cryosections from APAP-challenged RAG2^-/-^ mice treated with 150 μL of WT or RAG2^-/-^ serum. Gray: fibrin(ogen), red: IgM, green: IgG. Scale bar = 100 μm. (**B**) Quantification of the fibrin(ogen)^+^ area fraction in liver cryosections. (**C**) Serum ALT levels at 24 and 48 hours after APAP administration. (**D**) Quantification of Ki67^+^ hepatocytes in liver cryosections of RAG2^-/-^ mice after serum transfer. (**E**) Quantification of Ki67^+^ hepatocytes in liver cryosections of RAG2^-/-^ mice treated with 100 μg (i.v.) of purified IgM and IgG. See also figure S5G for Ki67 staining. Data are from RAG2^-/-^ mice that received an intravenous injection of WT serum or RAG2^-/-^ serum (**A-D**) or 100 μg purified IgG and IgM (**E**)4 hours after an oral gavage with 600 mg/kg APAP and were sacrificed 24 and 48 hours after the APAP challenge. Each dot in the graph represents a single mouse (**B-E**). Image quantifications were pooled from 10 fields of view (**B, D, E**). *p< 0.05.

However, RAG2^-/-^ mice injected with WT serum displayed an improved liver recovery 48h after injury, with significantly less hepatic necrotic areas than mice treated with RAG2^-/-^ serum (**Figures 5A and 5B**). Interestingly, the transfer of WT sera to RAG2^-/-^ mice was sufficient to increase the proliferation of hepatic cells 48h after injury, which was significantly elevated in comparison to RAG2^-/-^ mice that received RAG2^-/-^ serum (**Figure 5D and Supplementary 5G**). To confirm that the improvement of liver regeneration by NAbs restitution was not due to the presence of other serum components, RAG2^-/-^mice were treated with total IgM and IgG antibodies purified from WT sera. Similarly to the serum transfer experiment, treatment with purified antibodies alone significantly improved hepatocellular proliferation in the injured liver in comparison to RAG2^-/-^mice treated with vehicle (**Figure 5E**).

These data confirm that NAbs are central to drive liver repair after necrotic injury and that sera containing NAbs are sufficient to rescue the poor response to liver injury observed in RAG2^-/-^ mice.

### Natural antibodies drive the phagocytosis of necrotic cells debris through FcγRs and CD11b

As NAbs improved the recovery from necrotic liver injury, we further investigated the mechanism by which they exert their role in injury resolution. Antibody-opsonized antigens are known to be phagocytosed in an FcR-dependent manner, therefore, NAbs opsonization of necrotic cell debris likely promotes its clearance by phagocytosis. To test this hypothesis we developed an *in vitro* assay of necrotic debris phagocytosis by feeding necrotic cell debris to murine macrophage-like RAW 264.7 cells. Debris was prepared by crushing HepG2 cells and labeling it covalently with pHrodo Red succinimidyl ester, a pH-sensitive dye that emits increased fluorescence in acidified compartments such as phagosomes. This approach allowed us to discern the debris that was phagocytosed and matured in phagosomes from debris bound to or in proximity to cells. Labeled necrotic debris was left non-opsonized (PBS) or opsonized with mouse serum or with heat-inactivated (HI) serum (lacking complement activation). Opsonization with serum led to a significant increase in the phagocytosis of necrotic debris by RAW cells compared to non-opsonized (PBS) control samples (**Supplementary figure 6A**). Remarkably, necrotic debris phagocytosis was essentially identical if debris were opsonized with native or heat-inactivated serum, indicating that the complement cascade was not required in these conditions. In addition, incubation of RAW cells with latrunculin B, an inhibitor of actin polymerization known to prevent phagocytosis^31^, completely blocked the internalization of necrotic debris. We then investigated the FcRs expressed by RAW cells and found that they expressed constitutively the genes CD64 (FcγRI), CD32 (FcγRII), CD16 (FcγRIII) and, at lower level, CD16a (FcγRIV). However, the expression of the two FcRs that recognize IgM-coated targets, CD351 (Fcα/μR) and Faim3 (FcμR) was absent (**Supplementary figure 6B**). Stimulation of RAW cells with NAbs-opsonized necrotic debris for 6h increased the expression of CD64 and CD16a, but reduced the expression of CD32 (**Supplementary figure 6C-F**). Even after stimulation with NAbs-opsonized debris, we could not detect mRNA for CD351 and Faim3 in RAW cells (data not shown). The phagocytosis of NAbs-coated debris increased the expression of IL-10 while it did not alter the expression of TNF-α, suggesting an anti-inflammatory role of NAbs-mediated phagocytosis (**Supplementary figure 6G and 6H**).

After validating our methodology, we investigated necrotic debris phagocytosis in primary mouse neutrophils. Similar to our findings with RAW cells, serum-opsonized debris were efficiently phagocytosed by murine neutrophils and heat inactivation of serum did not alter the phagocytosis rate (**Figure 6A**), suggesting that NAbs but not complement is required in necrotic debris clearance. Importantly, the internalization of serum-opsonized debris is also completely inhibited by latrunculin B, indicating that phagocytosis is indeed the main process of internalization (**Figure 6A**). Our next step was to identify the receptors involved in the phagocytosis of NAbs-debris immunocomplexes. First, we performed a general blockage of FcγRs by treating neutrophils with 100 μg/mL purified mouse IgG, as previously described^32^, and we found a 50% reduction in phagocytosis efficiency of debris opsonized with normal serum or HI serum, suggesting that FcγRs and natural IgG are driving at least half of necrotic debris phagocytosis (**Figure 6A**). Considering that mouse neutrophils do not express Fc receptors for IgM^33,34^, we hypothesized that CD11b could have a role in the phagocytosis of IgM-coated debris, since CR3 has been implicated in the internalization of IgM/IgA-opsonized targets^35^. Thus, we performed a combined blockage of FcγRs and CD11b, which led to even further reduction of phagocytosis of serum-opsonized debris. Conversely, CD11b blockage did not alter the phagocytosis rate of HI-opsonized debris phagocytosis (**Figure 6A**).

**Figure 6:**
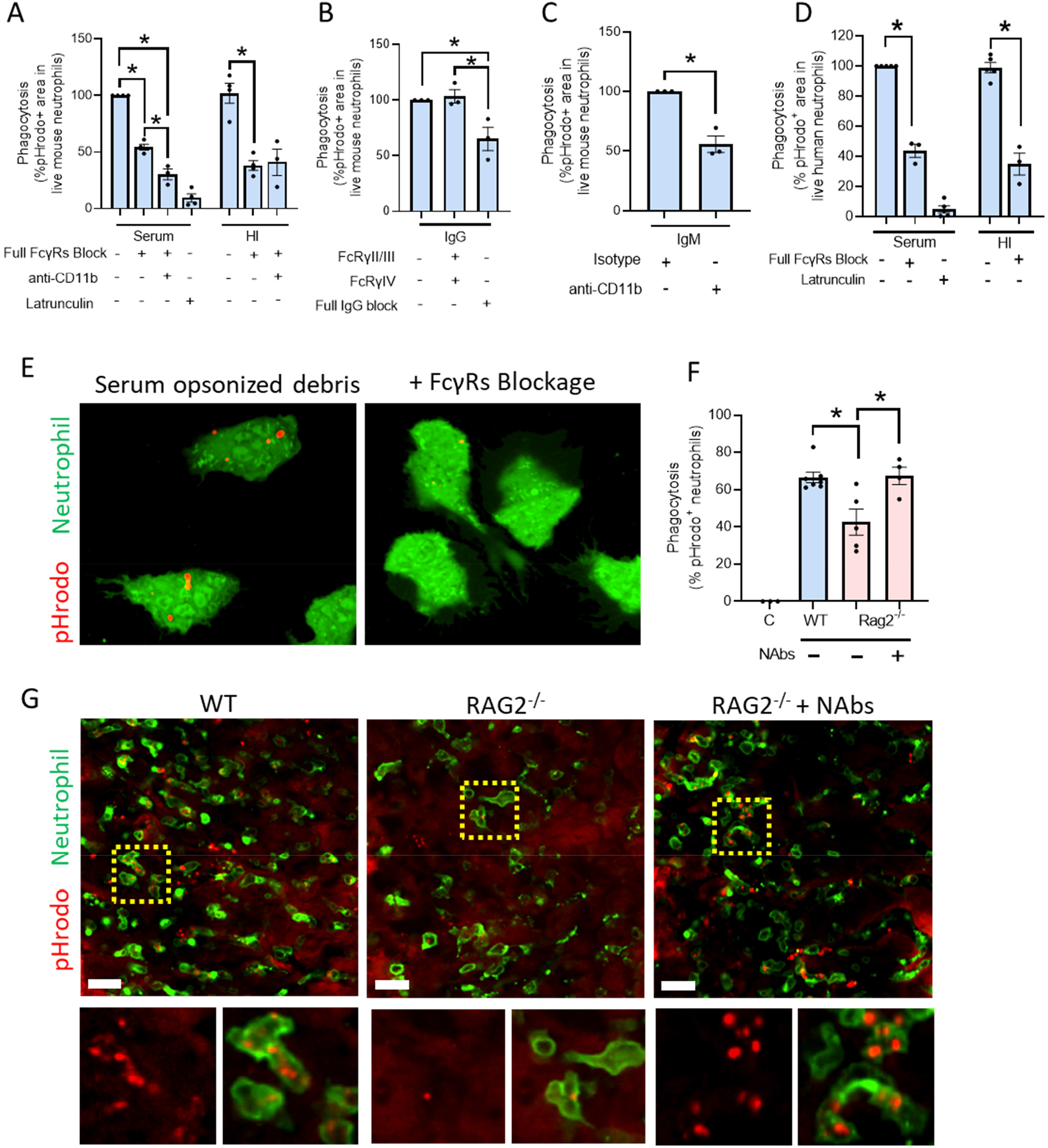
Natural antibodies drive the phagocytosis of necrotic cell debris through FcγRs and CD11b. (**A**) Phagocytosis by primary mouse neutrophils 3 hours after adding debris opsonized with normal or heat inactivated (HI) serum. FcγRs were blocked by adding 100 μg/mL IgG to the cells. CD11b was blocked with 10 μg/mL anti-mouse CD11b. Phagocytosis was blocked with 10 μM Latrunculin B. (**B and C**) Phagocytosis by primary mouse neutrophils 3 hours after adding IgG- or IgM-opsonized debris. Receptors were blocked using 10 μg/ml anti-CD16.2, anti-CD16/CD32, anti-CD11b or 100 μg/ml purified mouse IgG. (**D**) Phagocytosis by primary human neutrophils 3 hours after adding debris opsonized with normal or heat inactivated serum. FcγRs were blocked with 100 μg/mL IgG and Latrunculin B was used at 10 μM. (**E**) Representative confocal images from human neutrophils containing pHrodo^+^ phagosomes. FcγRs were blocked with 100 μg/mL purified human IgG. Scale bar = 10 μm. See also supplementary video 6. (**F**) Quantification of the percentage of neutrophils (Ly6G^+^, green) phagocytosing necrotic debris 6 hours after a focal burn injury in the liver of WT, RAG2^-/-^ mice. A separate group of RAG2^-/-^ mice were restituted with total purified Abs (100 μg/mouse). The focal burn was labeled with a droplet 4 μM pHrodo Red succinimidyl ester. Each dot in the graph represents a mouse. (**G**) Representative intravital microscopy images showing neutrophils (Ly6G^+^) phagocytosing necrotic debris as in F. Scale bar = 20 μm. Yellow square represents the area zoomed. Also see supplementary videos 3-5.

To better dissect the role of each immunoglobulin isotype, as well as to identify the FcγR involved in the recognition of IgG-coated debris, we performed the phagocytosis assay using necrotic debris opsonized with purified antibodies. Similar to our findings with serum-opsonized debris, the blockage of FcγRs using excessive IgG led to a significant reduction in the phagocytosis of IgG-coated debris. Surprisingly, the combined blockage of FcγRII, FcγRIII and FcγRIV did not affect phagocytosis, indicating that FcγRI might be the predominant phagocytic receptor for IgG-coated debris (**Figure 6B**). Using purified IgM-coated necrotic debris, CD11b blockage also resulted in a reduction of approximately 50% of phagocytosis, supporting a role for this integrin in the clearance of IgM-coated debris (**Figure 6C**). These data suggest that FcγRs are driving IgG-mediated phagocytosis of necrotic debris, whereas IgM-mediated phagocytosis is occurring at least in part via CD11b.

To investigate the mechanisms of NAbs-dependent necrotic debris phagocytosis in a more translational manner, we also used blood-derived human neutrophils. Primary neutrophils were also proficient in the phagocytosis of necrotic debris opsonized with native serum, a process that was once more prevented by latrunculin B (**Figure 6D**). Of note, no differences were observed in phagocytosis efficiency if necrotic debris was opsonized with donor-matched serum or not, corroborating the notion that NAbs share broad recognition of conserved molecules between individuals (data not shown). The phagocytosis of necrotic debris, quantified as the percentage of neutrophils containing pHrodo-positive events, was quite similar between normal and heat-inactivated serum, confirming that the clearance of necrotic debris occurs independently of the complement cascade (**Figure 6D**). Similarly to murine neutrophils, FcγR blocking in human neutrophils reduced the phagocytosis of necrotic debris by more than 50%, an inhibitory effect that was equal when debris was opsonized with native or heat-inactivated serum (**Figure 6D-E**). Using 3D confocal imaging 3h after adding serum-opsonized debris, we observed that neutrophils carried multiple pHrodo^+^ phagosomes and confirmed that the positive events were inside the cells (**Figure 6E and Supplementary video 3**). This indicates that, similarly to our data on mouse cells, the phagocytosis of necrotic debris by human neutrophils is complement-independent, while FcγRs drive a substantial fraction of the phagocytosis. Blockage of CD11b in human neutrophils, however, had no effect on the phagocytosis level (**Supplementary figure 6I**). Altogether, these findings show that the mechanisms of NAbs-mediated necrotic debris phagocytosis are shared between murine and human leukocytes. We also demonstrated that the phagocytosis of necrotic debris requires IgG and IgM NAbs and depends largely on FcγRs expressed on leukocytes.

### Natural antibodies are required for optimal debris phagocytosis at sites of necrotic injury *in vivo*

Considering the role of NAbs in the phagocytosis of necrotic cells and the defective recovery response observed in immunodeficient mice after liver injury, we sought to investigate if the absence of NAbs in these mice interfered with necrotic debris clearance in the liver. The experiments involved IVM imaging of a focal burn injury in the liver in which the whole necrotic area was labelled with a droplet of pHrodo Red succinimidyl ester (**Supplementary figure 6J**). After 6h, mice were injected i.v. with AF488-labeled anti-Ly6G to identify neutrophils migrating in the focal injury site. Approximately 66% of neutrophils in the injury site in WT mice contained pHrodo^+^ phagosomes, whereas RAG2^-/-^ mice presented a significant reduction in the number of pHrodo^+^ neutrophils (**Figure 6F and 6G**). To prove that the impaired phagocytosis of necrotic debris in RAG2^-/-^ mice was mainly due to the lack of NAbs, these mice were treated i.v. with 100 μg of purified IgM and 100 μg IgG 30 min prior to the focal burn injury. Restitution of NAbs to RAG2^-/-^ mice completely rescued the phagocytosis of necrotic debris by neutrophils to levels observed in WT mice (**Figure 6F, 6G and supplementary videos 4-6**). Importantly, no differences were found in the amount of recruited neutrophils inside necrotic areas between WT and RAG2^-/-^ mice (**Supplementary figure 6K**), and parameters of neutrophil migration including displacement, total distance, directionality, circularity and cell size were similar between the groups (**Supplementary figure 6L-Q**). These data suggest that NAbs are required for efficient clearance of necrotic cells debris by phagocytes *in vivo*, but not for leukocyte recruitment and migration in necrotic injury sites. In addition, it indicates that the delayed tissue repair in NAbs-deficient mice is correlated with impaired necrotic debris phagocytosis.

### Treatment with NAbs increases cellular proliferation and tissue regeneration after liver injury in immunocompetent mice

We evaluated whether the administration of NAbs could have protective effects also in normal mice. To assess this, WT mice were treated i.v. with a combination of 100 μg IgM and 100 μg IgG 4h after the APAP challenge and were evaluated 48h post injury. We observed that supplementation with NAbs significantly decreased the fibrin(ogen)^+^ necrotic areas in the liver (**Figure 7A and 7B**). Serum ALT levels were also reduced in the WT mice treated with NAbs (**Figure 7C**). Immunostaining of Ki67 in liver cryosections revealed that the treatment with NAbs significantly increased the hepatocellular proliferation 48h after APAP (**Figure 7D and 7F**). In addition, during necrotic injury the actin cytoskeleton labeling is lost within necrotic liver areas. However, as cells proliferate to regenerate the liver, the actin cytoskeleton scaffolds return, evidencing the repairing injury. By measuring the intensity of actin staining in centrilobular areas, we obtained further evidence that the treatment with NAbs improved significantly the regeneration of liver injury in WT mice (**Figure 7E and 7F**). In summary, treatment of immunocompetent mice with NAbs improved the resolution of injury and liver regeneration, suggesting that this approach could have therapeutic applications in immunocompetent individuals.

**Figure 7:**
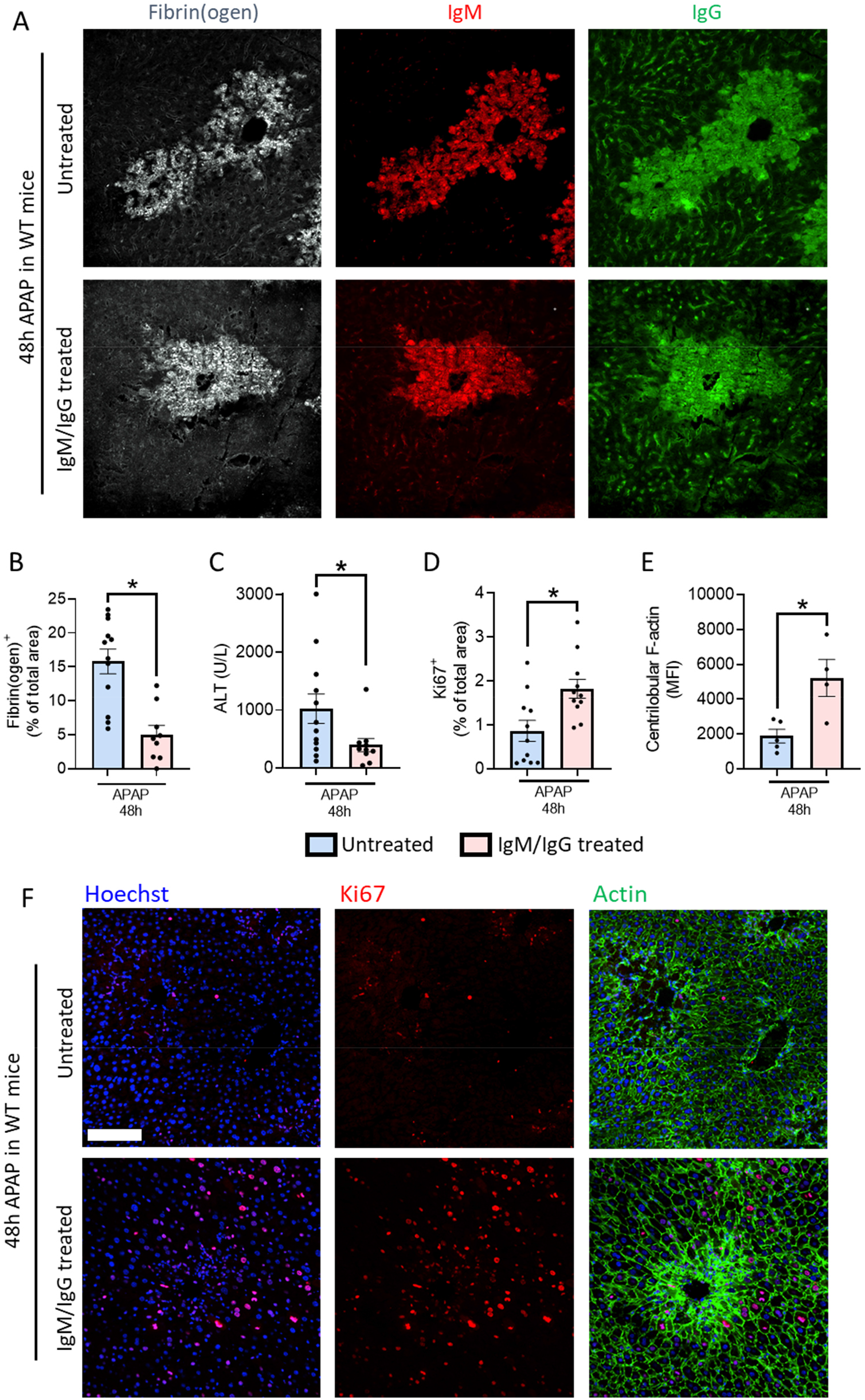
Treatment with NAbs increases cellular proliferation and tissue regeneration after liver injury in immunocompetent mice. (**A**) Representative images showing immunostaining of liver cryosections from WT mice treated with 100 μg/mouse of purified IgM and IgG, or isotype. Mice were treated 4h after the APAP challenge. Gray: fibrin(ogen), red: IgM, green: IgG. Scale bar = 100 μm. (**B**) Quantification of the fibrin(ogen)^+^ area in cryosections. (**C**) Serum ALT levels 48 hours after APAP administration. (**D**) Quantification of the Ki67^+^ area in WT mice treated with purified IgM and IgG or isotype (100 μg/mouse, i.v.) 48h after APAP challenge. (**E**) Mean fluorescence intensity of phalloidin staining around the centrilobular veins in WT mice treated with purified IgM and IgG or isotype (100 μg/mouse, i.v.) 48h after APAP. (**F**) Representative images of liver cryosections from WT mice showing parenchymal cell proliferation (Ki67^+^, red), F-actin (phalloidin, green) and nuclei (Hoechst, blue). Mice were treated with purified IgM and IgG (100 μg/mouse, i.v.). Scale bar = 100 μm. Data are represented as mean ± SEM. *p< 0.05.

## Discussion

Our work identified a so far unexplored physiological function of NAbs: to act as “eat-me” signals for the phagocytosis of necrotic cell debris in sites of tissue injury. We found that IgM and IgG NAbs rapidly bound necrotic debris and promoted their clearance partially via FcγRs and CD11b. NAbs-mediated clearance of necrotic cells increased hepatocellular proliferation and tissue recovery.

In mice, it is estimated that around 80% of serum IgM is natural IgM derived from B1 cells^36^. Its abundance, immediate availability, polyreactivity and targeting of conserved endogenous antigens makes NAbs ideal agents to clean up the disordered and heterogeneous necrotic cell debris. Here, we confirmed that NAbs opsonize diverse conserved molecules including DNA, actin, histones, phosphoinositides and cardiolipin, all of which are normally present only within cells. On the other hand, molecules that are found either in the outer leaflet of the plasma membrane or extracellularly such as PE, PC, cholesterol and triglycerides were not recognized by NAbs. A notable exception is PS, which although restricted to the plasma membrane inner leaflet in living cells, was not recognized by IgM or IgG NAbs. This is in accordance with previous work showing that the binding of natural IgM to apoptotic cells starts after these cells become positive for propidium iodide staining, suggesting that natural IgM only binds to late apoptotic/necrotic cells^5^. These observations indicate that Nabs recognize preferentially necrotic cells instead of apoptotic bodies and suggests that the immune system has evolved different mechanisms to deal with distinct types of cellular remnants.

The role of natural IgM in sterile liver injury has been investigated in previous works and provided unclear results. Marshall et. al. suggested that natural IgM participates in both the induction of liver ischemia-reperfusion injury and liver regeneration after 70% hepatectomy^22^. They evaluated two monoclonal IgM antibodies: B4, against annexin IV^37^ and C2, against a subset of phospholipids^38^. In our work, however, restitution of NAbs to immunodeficient mice did not increase injury. We believe this difference stems from our approach of reconstituting mice with the entire natural antibody repertoire, including natural IgG which was disregarded in previous works. Nevertheless, in accordance with their data, we also observed NAbs-dependent hepatocyte proliferation, which can be explained by several mechanisms: First, NAbs-dependent removal of debris will clear room for new cells to proliferate and reconstitute the dead parenchyma; Second, NAbs-dependent phagocytosis can promote the release of pro-resolutive mediators by phagocytes; Third, natural IgM and IgG can activate the complement cascade in the liver. Liver recovery is dependent on the activation of complement, as shown by impaired hepatic regeneration in C3-deficient mice, which was restored by C3a administration^39^. In C5-deficient mice, hepatocytes showed a marked delay to enter into the cell cycle (S phase) and reduced mitotic activity^40^. Thus, the mechanisms by which NAbs are beneficial may be multiple and synergistic, including liver regeneration but also resolution of inflammation.

Surprisingly, heat-inactivation of serum did not impact the efficiency of phagocytosis, suggesting that clearance of necrotic debris is independent of complement. We believe that this is due to multiple reasons, the first is that NAbs are sufficient to drive debris phagocytosis, thus, the absence of complement activation is easily overcome *in vitro.* Second, other phagocytic pathways will likely occur simultaneously, such as PS-dependent phagocytosis of necrotic cells^41^. Lastly, complement may require a higher concentration to opsonize necrotic debris effectively. By opsonizing debris in 20% serum, the role of the complement cascade may have been underestimated. Nevertheless, it has been shown that complement-mediated phagocytosis induces upregulation of genes encoding proinflammatory cytokines in mouse macrophages such as TNF-α^42^, while our NAbs-dependent phagocytosis of necrotic debris did not alter TNF-α expression but upregulated IL-10.

We showed that the phagocytosis of necrotic debris immunocomplexes is largely dependent of FcγRs, since blockage of these receptors in neutrophils reduced phagocytosis by 50%. Dual blockage of FcγRs and CD11b decreased phagocytosis even further (70%), suggesting also a role for CD11b likely via IgM recognition. However, CD11b blockage did not affect the phagocytosis of HI-opsonized necrotic debris. We believe that this is due to the heat inactivation process also affecting the polygonal structure of IgM, thereby reducing its binding to antigen and recognition by receptors. In fact, recent work suggested that heat-inactivation reduces IgM values in serum in a temperature-dependent manner^43^. Moreover, a well characterized method to produce multimeric IgG is by heat-induced aggregation^44^, which increases both the avidity to antigens and recognition by FcRs in macrophages^45–47^. Thus, heat-inactivation may favor the role of IgG in promoting phagocytosis *in vitro*.

Although the phagocytosis of necrotic debris is mainly driven by NAbs, the remaining phagocytosis can be occurring via other opsonins, e.g. PS-binding proteins and calreticulin. *In vivo*, the reduction of phagocytosis in RAG2^-/-^ was approximately 40%. Especially in these experiments, it is most likely that RAG2^-/-^ develop compensatory mechanisms for the lack of NAbs. Since the complement cascade was shown to be activated during several types of liver injury, it is quite possible that the complement cascade contributes to the clearance of necrotic debris observed in immunodeficient mice. However, to test the combined deficiency in both the complement cascade and NAbs in necrotic debris clearance may be proven impossible, since it is unlikely that a deficiency in two of the most central components of the immune system would be tolerable.

Altogether, our study provides a new insight on the clearance of necrotic remnants from tissues, highlighting NAbs as adaptors for the phagocytosis of these potentially dangerous self-antigens. Considering that phagocytes have a key role in the clearance of necrotic debris, immunosuppressants and therapies that inhibit leukocyte recruitment must be carefully evaluated. Conversely, a therapy based on NAbs may arise as a pro-resolutive alternative by increasing debris clearance and tissue repair, rather than preventing inflammation from occurring.

### Limitations of the study

While our study demonstrates that NAbs are required for efficient phagocytosis of necrotic debris in mice and in human cells *in vitro*, it remains to be evaluated if NAbs are as important in human physiology as they are in mice. Here, we used two *in vivo* models of necrotic liver injury (APAP-induced liver injury and focal burn in the liver), however, it should be determined if NAbs are relevant in other diseases as well as in different organs. The reader should consider that differences in tissue composition and vascularization may be important in this mechanism. The liver is heavily vascularized and poor in extracellular matrix in comparison with skin and adipose tissue, for example. The intimate contact of hepatic cells with the bloodstream may be determinant for the extensive opsonization by NAbs that was observed. Investigation of the role of NAbs in different acute injury models is warranted especially in the skin, since it is a tissue so often exposed to damage. Another relevant point would be to discern the individual impact of IgM and IgG in the clearance of necrotic debris and the consequent inflammatory response to injury: do they favor resolution? Are they coupled to the activity of pro-resolving mediators? Non-phlogistic phagocytosis? Studies following on these questions could increase the potential for therapies that increase the clearance of necrotic debris in several human diseases.

## RESOURCE AVAILABILITY

### Lead contact

Further information and requests for resources and reagents should be directed to and will be fulfilled by the lead contact, Pedro Elias Marques.

### Material availability

This study did not generate new unique reagents.

### Data and code availability

Any additional information required to reanalyze the data reported here, is available from the lead contact upon request.

## EXPERIMENTAL MODEL AND SUBJECT DETAILS

### Mice

C57BL/6J and C57BL/6NRj mice were purchased from Janvier Labs. C57BL/6N-Rag2Tm1/CipheRj (RAG2^-/-^) were bred in specific pathogen-free conditions at the Animal Facility of the Rega Institute (KU Leuven). C57BL/6-Ly 5.1 and IgMi mice were provided by Ari Waisman. All mice used in this study were between 10-12 weeks old and both male and female mice were equally distributed across experiments (no phenotypic differences between genders were observed). Mice were housed in acrylic filtertop cages (5 mice per cage) with an enriched environment (bedding, toys and small houses), at the Animal Facility of the Rega institute (KU Leuven). Water and food were provided *ad libitum* and mice were kept under a controlled dark/light cycle (12/12 h) at 21 °C. All experiments were approved and performed following the guidelines of the Animal Ethics Committee from KU Leuven (registry number: P125/2019).

### Cell Lines

RAW 264.7 is a monocyte/macrophage like cell that was derived from a tumor induced in a male mouse with the Albeston murine leukemia virus. HepG2 cell is a hepatocyte-like cell derived from a hepatocellular carcinoma of a 15-year-old male human. The cells were cultured at 37 °C and 5% CO2 atmosphere in high glucose Dulbecco’s Modified Eagle Medium (DMEM) with GlutaMAX (Thermo Fisher Scientific), supplemented with 10% FBS (Sigma-Aldrich), 1 mM sodium pyruvate (Thermo Fisher Scientific) and 0.12% sodium bicarbonate (Thermo Fisher Scientific).

## METHOD DETAILS

### Acetaminophen (APAP)-induced liver injury model

Mice were fasted for 15 hours before a single oral gavage of vehicle or APAP (600 mg/kg, Sigma-Aldrich, St. Louis, MO, USA) dissolved in warm PBS. Fasting was performed to guarantee full APAP absorption and to increase the reproducibility amongst the experiments. After 6, 12, 24, 48 or 72 hours, mice were sacrificed under anesthesia containing ketamine (80 mg/kg) and xylazine (4 mg/kg) whereafter liver and blood were harvested.

### Alanine aminotransferase (ALT) assay

Liver injury was indirectly assessed by monitoring levels of serum ALT utilizing a kinetic test (Infinity, Thermo Fisher Scientific, Waltham, MA, USA). Briefly, blood samples were harvested and centrifuged for 10 minutes at 1500 x g and then serum was harvested. Pure serum and three different dilutions (1:10, 1:20, 1:30) were added to a 96-well plate, then, the substrate (HEPES buffer pH 7.8, LDH, L-Alanine, NaCl) and coenzymes (alfacetoglutarate and NADH) were added to the serum samples at 37 °C. The reaction was monitored every minute (for a total of 3 minutes) by measuring the rate of decrease in absorbance at 340 nm (CLARIOstar, BMG Labtech, Cary, NC) due to the oxidation of NADH to NAD.

### Liver cryosectioning and immunostaining

To perform liver immunostainings, the left lobe was harvested, embedded in frozen mounting medium (PolyFreeze, Sigma-Aldrich, St. Louis, MO) and snap-frozen in liquid nitrogen. Cryosections of 14 μm thickness were generated using a cryostat (Microm Cryo-Star HM560, Thermo Fisher Scientific, Waltham, MA, USA). The sections were fixed for 1 hour with 4% paraformaldehyde (PFA) in Hank’s Balanced Salt Solution (HBSS, Gibco, Waltham, MA, pH 7.2) supplemented with 0.1% Bovine Serum Albumin (BSA, Albumin Fraction V, protease-free, Carls Roth, Karlsruhe, DE) at room temperature (RT). Then, the sections were washed with HBSS and permeabilized with 0.1% Triton X-100 in HBSS for 1 hour at RT. Livers sections were washed again and blocked using 1% Fc Block (MACS, Miltenyi Biotec, Auburn, CA, USA) and 5% BSA in HBSS during 1 hour at RT. To visualize the necrotic areas in liver cryosections, we stained for fibrin(ogen), once the deposition of fibrin(ogen) within areas of hepatocellular necrosis is well described^48,49^. After another washing step, the primary polyclonal rabbit anti-human/mouse fibrin(ogen) (10 μg/mL in HBSS, Agilent Dako, Glostrup, DK) antibody was added overnight at 4°C. The sections were washed and the secondary antibodies Alexa Fluor 488 donkey anti-rabbit, Rhodamine RED-X (RRX) donkey anti-mouse IgM and Alexa Fluor 647 goat anti-mouse IgG (all at 10 μg/mL, Jackson ImmunoResearch, West Grove, PA, USA) were added for 3 hours at RT. After another washing step, mounting medium was applied (ProLong Diamond, Thermo Fisher Scientific) and the sections were imaged using an Andor Dragonfly 200 spinning-disk confocal microscope equipped with a 25X objective, and analyzed using FIJI.

### Purification of liver non-parenchymal cells (NPCs)

The purification of liver NPCs was performed as previously described^26^. Briefly, the caudate and median liver lobes were harvested in RPMI-1640 medium (Biowest Riverside, MO, US) and mechanically minced using the gentleMACS Dissociator (Miltenyi Biotec, Auburn, CA, USA). Then, RPMI-1640 medium was added to the liver homogenate to complete 30 ml and centrifuged at 300 x g for 5 minutes at 4 °C. The supernatant was discarded and more RPMI-1640 medium was added to complete 30 ml. After homogenization, the cells were centrifuged at 60 x g for 3 minutes at 4 °C and the supernatant, containing the NPCs, was harvested and filtered through a 40 μm cell strainer in order to remove undigested tissue. The cells were centrifuged again at 300 x g for 5 minutes at 4 °C and the supernatant was discarded. The pellet was resuspended, transferred to a 15 mL tube and centrifuged again (300 xg, 5 minutes at 4 °C). The supernatant was discarded and red blood cells were lysed with 2 ml of ACK Lysing Buffer (Gibco, Grand Island, NY, US) for 5 minutes on ice. ACK was washed away by adding PBS until 10 ml and centrifuged at 300 x g for 5 minutes at 4 °C. The final pellet, containing the liver NPCs, was reconstituted for further analyses.

### Flow Cytometry

For flow cytometry, 5 × 10^5^ NPCs were labeled with viability dye marker (Zombie Aqua, Biolegend, San Diego, CA, USA) and blocked with PBS supplemented with 1% Fc Block (MACS, Miltenyi Biotec) and 0.5% BSA for 15 minutes at 4 °C. The cells were washed with 500 μL FACS buffer (PBS supplemented with 0.5% BSA and 2mM EDTA) and labeled with different antibodies (see Key resources Table) for 30 minutes at 4 °C in the dark. After labeling, the samples were washed by adding 500 μL FACS buffer, centrifuged at 300 x g for 5 min at 4 °C and resuspended in 300 μl FACS buffer. Cells were immediately read in a Fortessa X20 (BD Biosciences, Franklin Lakes, NJ US). The data was analyzed using FlowJo X (FlowJo 10.8.0, LLC, Asland, OR USA). The gate strategy can be seen at supplementary figure 7.

### Hepatocyte debris spot

Livers from healthy mice were harvested in RPMI-1640 medium (Biowest Riverside, MO, US) and mechanically minced using a gentleMACS Dissociator. Then, RPMI-1640 medium was added to the liver homogenate to complete 30 ml and centrifuged at 300 x g for 5 minutes at 4 °C. The supernatant was discarded and more RPMI-1640 medium was added to complete 30 ml and centrifuged at 60 x g for 3 min at 4 °C. The supernatant was discarded and the liver homogenate was transferred to a new 50 mL tube and filtered through a 40 μm cell strainer in order to remove the undigested tissue. Samples were centrifuged again at 300 x g for 5 minutes at 4 °C. The supernatant was discarded and red blood cells were lysed by adding ACK to the pellet (containing the hepatocytes) for 5 minutes on ice. The cells were then washed with HBSS and centrifuged at 300 x g for 5 min at 4 °C. 1 × 10^7^ hepatocytes were mechanically disrupted with a pellet mixer (VWR Radnor, PA, USA) for 5 minutes. After crushing the cells, the debris spot was generated by adding 6 μL of the solution containing the necrotic hepatocytes into an 8-well chamber slide (Nunc, Rochester, NY, USA) and drying for 2 hours in the laminar flow. The hepatocyte debris spot was then blocked with HBSS supplemented with 1% Fc Block and 0.5% BSA for 15 min at RT. After washing three times with HBSS, healthy mouse serum (1:10 in HBSS) was added for 30 min at 37 °C. The hepatocyte debris spots were then washed three times and stained with Hoechst (10 μg/mL), Phalloidin (66 nM) and secondary antibodies Rhodamine RED-X (RRX) rabbit anti-mouse IgM and Alexa Fluor 647 goat anti-mouse IgG (both 10 μg/mL, Jackson ImmunoResearch, West Grove, PA, USA) for 1 hour at RT. After another washing step, the hepatocytes debris spots were imaged using a Zeiss Axiovert200M microscope and analyzed with FIJI. Images were taken using an 25X objective.

### Measurement of natural antibodies in the serum

To assess the levels of circulating NAbs during APAP-induced liver injury, serum was collected from healthy mice and from mice 6, 12, 24, 48 and 72 hours after APAP administration. Then, hepatocyte debris spots were incubated with the serum for 30 min at 37 °C. The hepatocyte debris spots were then washed three times and stained with Hoechst (10 μg/mL), phalloidin (66 nM) and secondaries antibodies Rhodamine RED-X (RRX) rabbit anti-mouse IgM and Alexa Fluor 647 goat anti-mouse IgG (both 10 μg/mL, Jackson ImmunoResearch, West Grove, PA, USA) for 1 hour at RT. After another washing step, the hepatocyte debris spots were imaged using a Zeiss Axiovert200M microscope and the mean fluorescence intensity (MFI) of IgM and IgG were determined using FIJI. The data are represented as the relative change in the MFI compared to the control (healthy mouse serum) group.

### Serum adoptive transfer and treatment with purified IgG and IgM

For the serum adoptive transfer, WT or RAG2^-/-^ mice were euthanized under anesthesia and blood was harvested. Serum was collected by centrifugationat 1500 x g for 10 min at 4 °C. Next, RAG2^-/-^mice were injected with either 150 μL of WT serum or with the same amount of RAG2^-/-^ serum. For the purified antibodies treatment, mice received 100 μg of purified IgG and 100 μg of purified IgM diluted in sterile PBS in a final volume of 200 μL. As control for the purified antibodies injection, control mice received an injection with the same amount of IgG and IgM monoclonal isotype control (see key resources table). The serum or purified antibodies were administered intravenously, only once, through the retro-orbital sinus 4 hours after APAP challenge. During the injection, mice were anaesthetized with isoflurane.

### Dot blot and lipid blot assays

For the dot blot assay, purified DNA (Sigma-Aldrich), actin (Sigma-Aldrich) and histones (Cytoskeleton, Inc.) were diluted in Milli-Q water to a final concentration of 2 mg/mL. Then, 2 or 4 μg of purified debris was spotted on nitrocellulose membranes and dried for 1 hour at RT. For the lipid blot, PIP Strips and Membrane Lipid Strips (Echelon Biosciences, Salt Lake City, UT, USA) containing different lipids at 100 pmol per spot were used. Unspecific labeling was blocked by adding washing buffer (Tris 20 mM, NaCl 150 mM and Tween 20 0.1%) supplemented with 5% BSA for 1 hour at RT under constant agitation. The membrane was washed three times for 5 minutes with washing buffer. After washing, the membrane was incubated with mouse serum (1:10 in washing buffer) or 5 μg/mL of purified IgG and IgM (Rockland, Limerick, PA, USA) in washing buffer overnight at 4°C under agitation. The membrane was washed again three times for 5 minutes each, and incubated with the secondary goat anti-mouse IgM IRDye 680RD and goat anti-mouse IgG IRDye 800CW (1:10,000, LI-COR Biosciences, Lincoln, NE, USA) for 3 hours in the dark at RT with agitation. The membrane was washed again as aforementioned and imaged with an Odyssey Fc Imaging System (LI-COR Biosciences) at 800 and 700 nm. The images were analyzed with Image Studio lite (LI-COR Biosciences).

### Analysis of chemokines and cytokines

Levels of cytokines and chemokines were assessed in serum samples by sandwich ELISA using commercial kits (DuoSet R&D Systems, Minneapolis, MN, USA) following the manufacturer’s protocol. The absorbance was determined using a spectrophotometer (CLARIOstar, BMG Labtech, Cary, NC) at 490 nm. Results were represented as pg/mL of cytokines or chemokines.

### Neutrophil purification

Human neutrophils were purified from blood of healthy volunteers with the EasySep neutrophil isolation kit (StemCell Technologies, Vancouver, Canada) following the manufacturer’s instructions. Mouse bone marrow (BM) neutrophils were extracted from femurs and tibias of C57BL/6J mice by flushing the bones with 5 mL cold RPMI-1640 medium using a 26 gauge needle. Cells were filtered through a 70 μm nylon strainer and further purified with the EasySep mouse neutrophil enrichment kit (StemCell Technologies, Vancouver, Canada), following the manufacturer’s instructions.

### Human IgG purification

10 ml of human plasma from 4 healthy volunteers was pooled, filtered (0.45 μm) and diluted 1:1 in PBS. A 5 ml protein G Sepharose 4 Fast flow column (GE Healthcare) was used to extract total human IgG. Sample was loaded (1 ml/minute) and washed with 4 column volumes PBS. Bound antibodies were eluted using 0.1 M glycine (pH 2.3), and 1 M Tris-HCl (pH 8.0) was used to neutralize the eluted fractions. The antibody concentration was determined by extinction coefficient with an absorbance at 280 nm of 1.4 being equal to a concentration of 1.0 mg of IgG.

### In vitro neutrophil phagocytosis assay

For the phagocytosis assay, purified human neutrophil or mouse BM-derived neutrophils were stimulated with 10^-7^M N-formyl-Met-Leu-Phe (fMLF; Sigma-Aldrich) or 1 μM WKYMV (Phoenix Pharmaceuticals, Germany), respectively, labeled with 1 μM calcein AM viability dye (Invitrogen) and seeded in a 48-well plate at 50 × 10^3^ cells per well. Receptors were blocked by adding 100 μg/ml purified human IgG, 100 μg/ml purified mouse IgG (Sigma-Aldrich), 10 μg/ml anti-mouse CD16.2 (BioCell), 10 μg/ml anti-mouse CD16/CD32 (BD Pharmingen), 10 μg/ml anti-human/mouse CD11b (M1/70 clone, Biolegend) antibody or 10 μg/ml purified rat IgG2b isotype (Biolegend), 10 minutes prior adding opsonized debris.

Necrotic debris was generated by mechanical disruption of HepG2 cells with a pellet mixer for 5 minutes. The debris was washed with PBS (5 min, 13000 x g, RT) and labeled for 1 hour with pHrodo Red succinimidyl ester (Thermo Fisher Scientific) with 2 μL of a 10 mM solution per 10×10^6^ cells, in 0.1 M sodium bicarbonate at pH 8.4. The unbound pHrodo was washed away (5 min, 13000 x g, RT), whereafter the debris was opsonized with 20% fresh mouse/human serum, mouse/human heat-inactivated serum (30 min, 56°C) in PBS, 10 μg/mL purified mouse IgG, or 10 μg/mL purified mouse IgM (Mouse IgG and IgM whole molecule, Rockland, Limerick, PA, USA) for 1h at 37°C. The debris was washed with PBS (5 min, 13000 x g, RT) and added to the neutrophils in a 1:10 (cells/debris) ratio. 10 μM latrunculin B, an actin polymerization inhibitor (Sigma-Aldrich), was used to block phagocytosis. To ensure that the necrotic debris would reach the cells in a fast and homogenous way, the plate was centrifuged for 5 min, 300 x g at 4°C. Images were taken every 30 min with the Incucyte Live-Cell Analysis (Sartorius).

### *In vitro* RAW cell phagocytosis assay

50 × 10^3^ RAW 264.7 cells were seeded in a 48-well plate (Corning) overnight at 37°C. Living RAW 264.7 cells were labeled with 1 μM calcein acetoxymethyl ester (AM) viability dye (Invitrogen) for 20 min at 37°C in FBS free medium. Necrotic debris was generated by mechanical disruption of HepG2 cells with a pellet mixer for 5 minutes. The debris was washed with PBS (5 min, 13 000 x g, RT) and labeled for 1 hour with pHrodo Red succinimidyl ester (Thermo Fisher Scientific) with 2 μL of a 10 mM stock per 10×10^6^ cells, in 0.1 M sodium bicarbonate at pH 8.4. The unbound pHrodo was washed away with PBS (5 min, 13 000 x g, RT) whereafter the debris was opsonized with 20% fresh mouse serum, heat-inactivated (30 min, 56°C) serum in PBS, 10 μg/mL purified IgG or 10 μg/mL purified IgM (Mouse IgG and IgM whole molecule, Rockland, Limerick, PA, USA) for 1h at 37°C. The debris was washed with PBS (5 min, 13 000 x g, RT) and added to the RAW cells in a 1:10 (cells/debris) ratio. 10 μM latrunculin B (Sigma-Aldrich) was used to block phagocytosis. After centrifugation (5 min, 300 x g, 4°C), images were taken every 30 min with the Incucyte Live-Cell Analysis (Sartorius).

### *In vivo* phagocytosis assay

Mice were anesthetized by a subcutaneous injection of 80 mg/kg ketamine and 4 mg/kg xylazine. Then, a small midline incision was made in the abdominal area to expose the liver. With a hot needle (26G), a liver burn injury of approximately 1 mm^3^ was made on which a droplet of pHrodo Red succinimidyl ester (4 μM; Thermo Fisher Scientific) was administered. The incision was stitched and after 6h mice were again anaesthetized with ketamine and xylazine for imaging of the burn site by intravital microscopy. For the restitution of NAbs, RAG2^-/-^ mice were treated with purified IgM and IgG antibodies (100 μg each) intravenously 30 minutes prior the focal burn injury.

### Confocal intravital microscopy

Intravital imaging was performed as previously described^26,50^. Mice were anesthetized by subcutaneous administration of 80 mg/kg ketamine and 4 mg/kg xylazine. Then a midline laparotomy was performed to expose the liver for imaging. 10 minutes before the surgery, mice received an intravenous injection of 100 μL of a mixture containing Sytox green (2 μL/mouse at 5mM, Thermo Fisher Scientific, Waltham, MA) and 200 μg/kg anti-mouse Ly6G Alexa Fluor 488 antibody (Biolegend) diluted in PBS. Specific labeling was confirmed by the injection of matched isotypes controls. Mice were imaged every 30 seconds using a 25X objective in an Andor Dragonfly 200 series high speed confocal platform system (Oxford Instruments, Abingdon, OX, UK).

### Image analysis and neutrophil tracking

Neutrophil tracking and morphology analysis were performed using the plugin TrackMate 7, in FIJI, following the developer’s methodology^51^. Neutrophils were detected using a thresholding detector that recognizes gray pixels that have a value greater than the established threshold. Linear Assignment Problem was used as a tracker adapted from Jaqaman et.al., 2008^52^. Tracking analysis was performed using at least four different animals per group.

### RNA extraction and qPCR

After the phagocytosis experiments, cells were harvested and total RNA was extracted using a Rneasy Plus Mini Kit (Qiagen, Hilden, Germany) following the manufacturer’s instructions. After extraction, total RNA was quantified using a Nanodrop and 1 μg of RNA was used to perform the reverse transcription using a High-Capacity cDNA Reverse Transcription Kit (Applied Biosystems, Waltham, MA, US). 50 ng of the resulting cDNA was amplified in a 7500 Real-Time PCR system (Applied Biosystems) using IDT primers (See key resource table) and the TaqMan Gene Expression Master Mix (Applied Biosystems). RT-qPCR data were expressed as 2^-ΔΔCT^ relative to gene expression of cells in steady-state. CT values were obtained using *Cdkn1a* as housekeeper gene.

## QUANTIFICATION AND STATISTICAL ANALYSIS

### Statistical analysis

All statistical analysis was performed in GraphPad Prism v9.3.1. Significance between two groups was analyzed with Student *t* test and between multiple groups with one-way ANOVA. Differences were considered significant if p ≤ 0.05. Grubb’s test (extreme studentized deviate) was applied in the data in order to determine whether extreme values were significant outliers from the rest. Data were represented as ± SEM.

## Acknowledgments

This work was supported by the Research Foundation of Flanders (FWO-Vlaanderen) Junior Research Grants (G058421N and G025923N). MSM was supported by an EILF-EASL Sheila Sherlock Post-graduate Fellowship. SV and SS hold PhD fellowships from FWO-Vlaanderen (SB1S56521N and 1116922N, respectively). PEM is supported by a Marie Sklodowska-Curie Fellowship (MSCA-IF-2018-839632) and the Rega Foundation. AW and NH were supported by Deutsche Forschungsgemeinschaft (DFG, German Research Foundation) project number 318346496 – SFB1292/2 and project number 490846870-TRR355/1.

## Author contributions

MSM and PEM designed the experiments. MSM, SV and PEM wrote the manuscript; MSM, SV, SS, LF and NH conducted the experiments; AW, MSM and PEM provided reagents, analyzed and discussed data.

## Inclusion and diversity

We support inclusive, diverse, and equitable conduct of research.

