## Supplemental figures for "Natural antibodies as “eat-me” signals for phagocytosis of necrotic cell debris at sites of tissue injury"

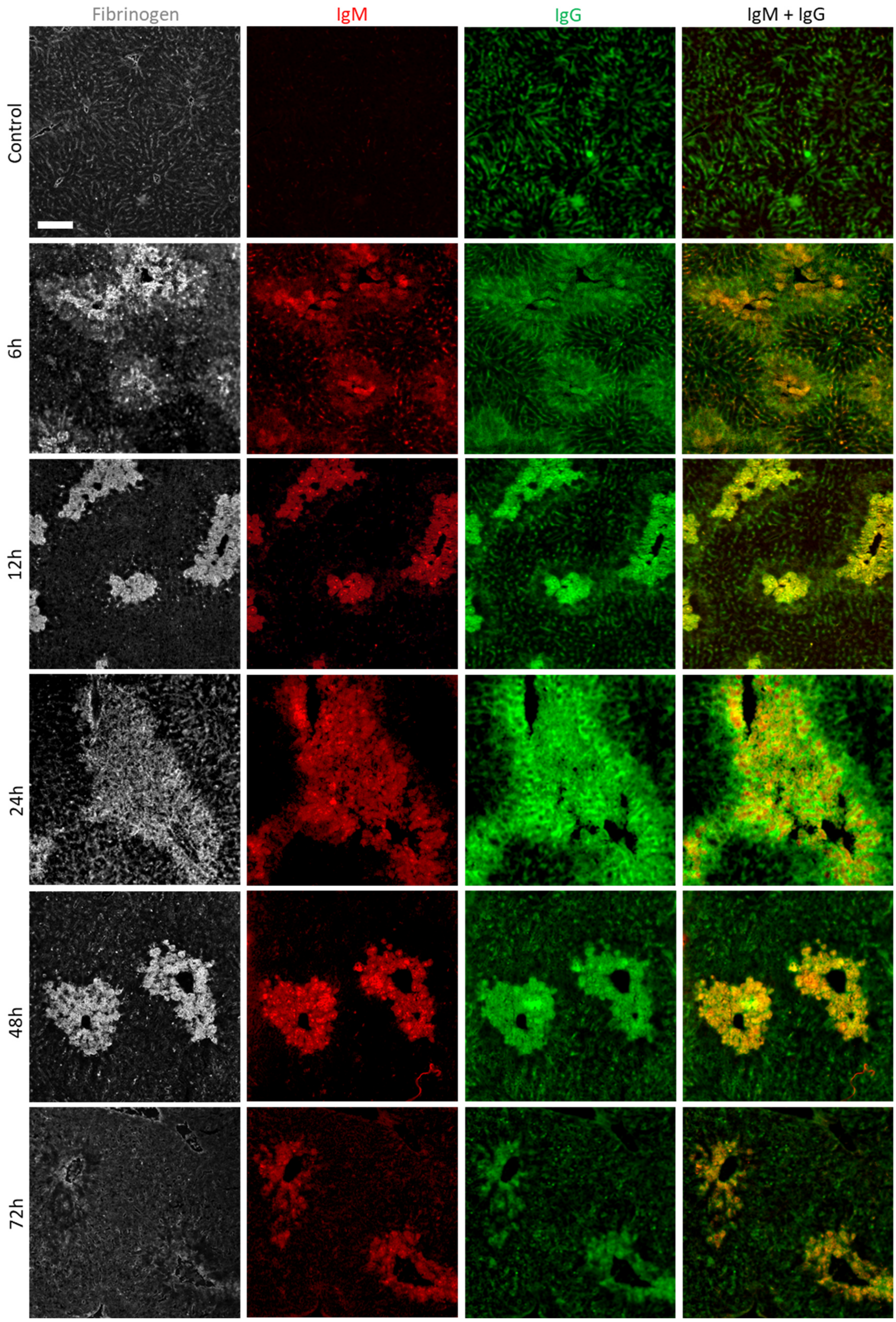


**Supplementary Figure 1:** (A) Representative images showing immunostaining of liver cryosections from control mice or mice gavaged with 600 mg/kg APAP after 6, 12, 24, 48 or 72 hours. Gray: fibrin(ogen), red: IgM, green: IgG, orange: merged IgM and IgG. Scale bar = 100 µm.

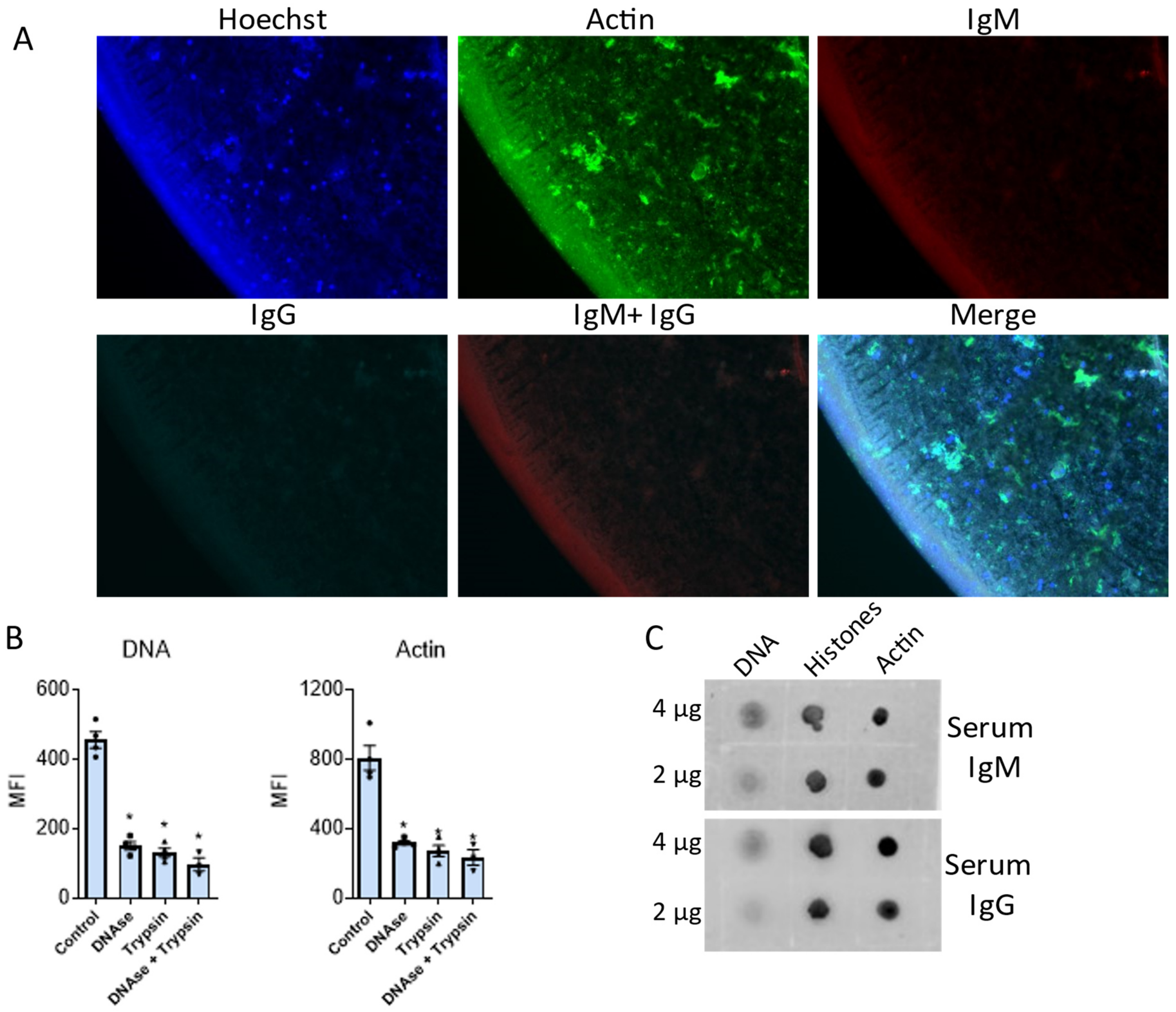


**Supplementary Figure 2:** (A) Representative images of a necrotic hepatocyte debris spot incubated with secondary anti-mouse IgM (red) and IgG (cyan) only. DNA is in blue (Hoechst) and f-actin (phalloidin) in green. (B) Mean fluorescence intensity (MFI) of DNA and actin labeling after pre-treatment of the debris spot for 15 minutes with DNase and/or Trypsin. Data are represented as mean ± SEM. *$p < 0.05$ compared to untreated samples. (C) Dot blot showing the reactivity of serum natural IgM and IgG to 2 µg or 4 µg of purified DNA, histones and actin.

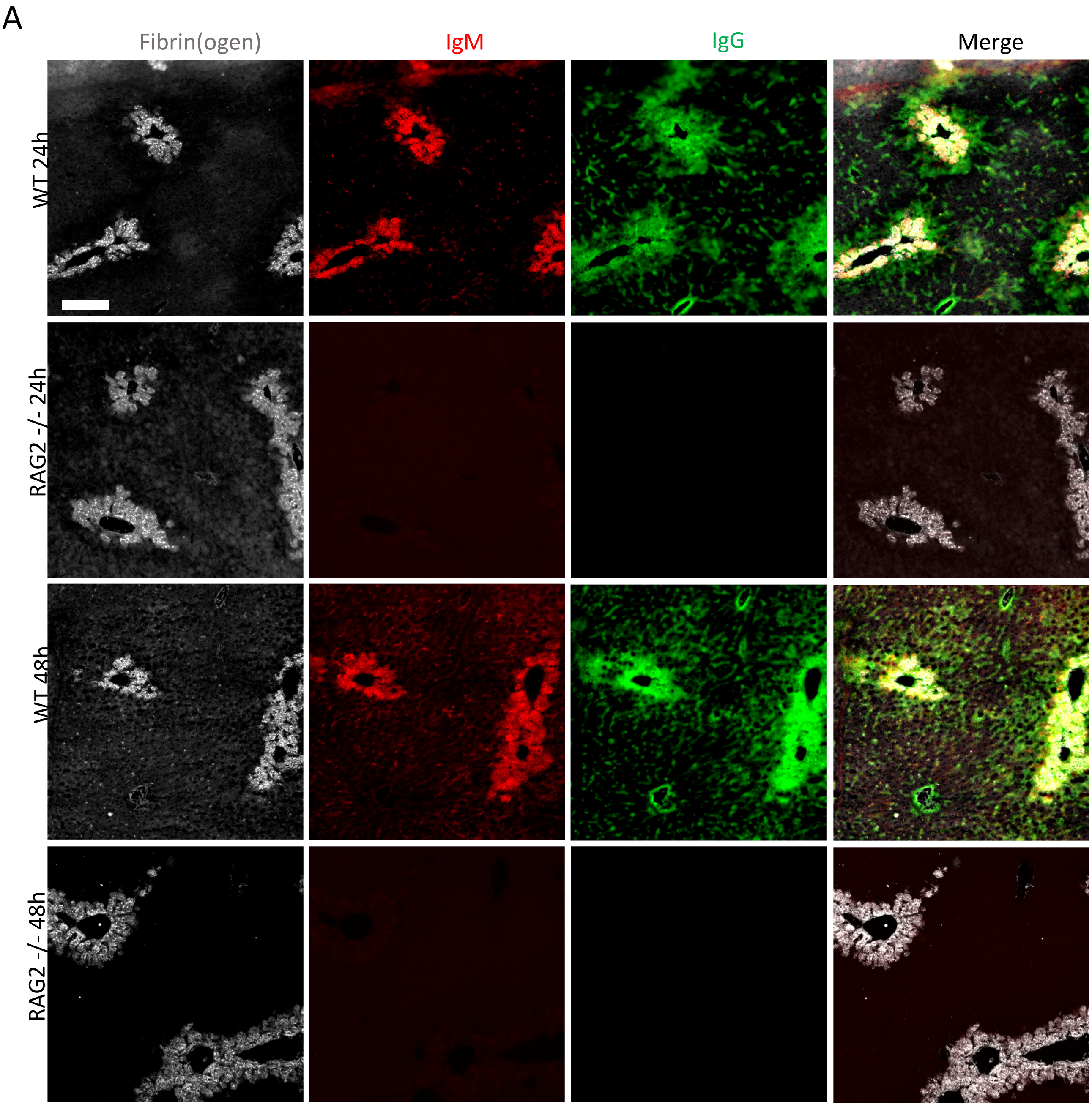


**Supplementary Figure 3:** (A) Representative immunostaining images of liver cryosections from WT and RAG2$^{-/-}$ mice, 24 hours and 48 hours after oral gavage with 600 mg/kg APAP. Gray: fibrin(ogen), red: IgM and green: IgG. Scale bar = 100 µm.

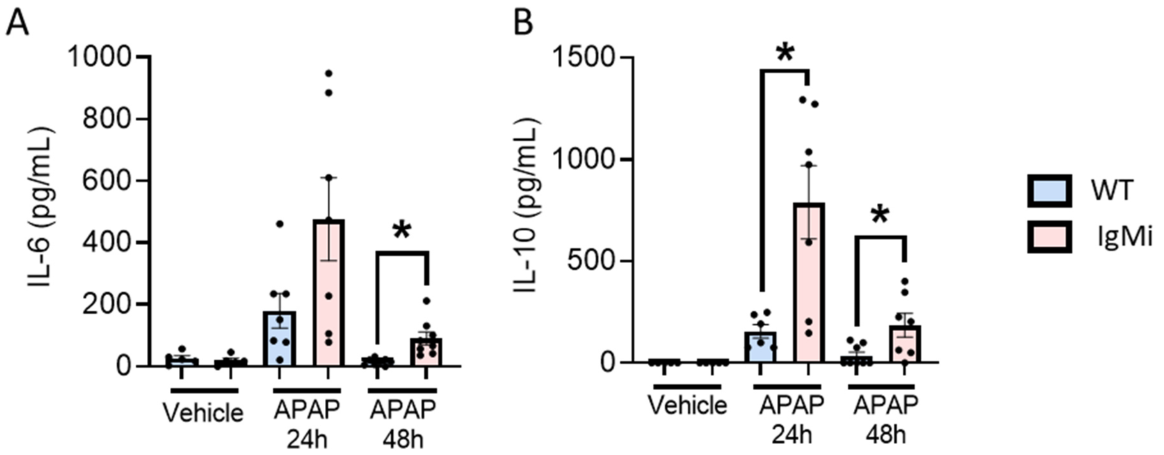


**Supplementary Figure 4:** (A) Serum levels of IL-6 in WT and IgMi mice after receiving APAP (600 mg/kg) for 24 and 48 hours. (B) Serum levels of IL-10 in WT and IgMi mice after receiving APAP (600 mg/kg) for 24 and 48 hours. Each dot in the graphs represents a single mouse. Data are represented as mean ± SEM. *$p < 0.05$.

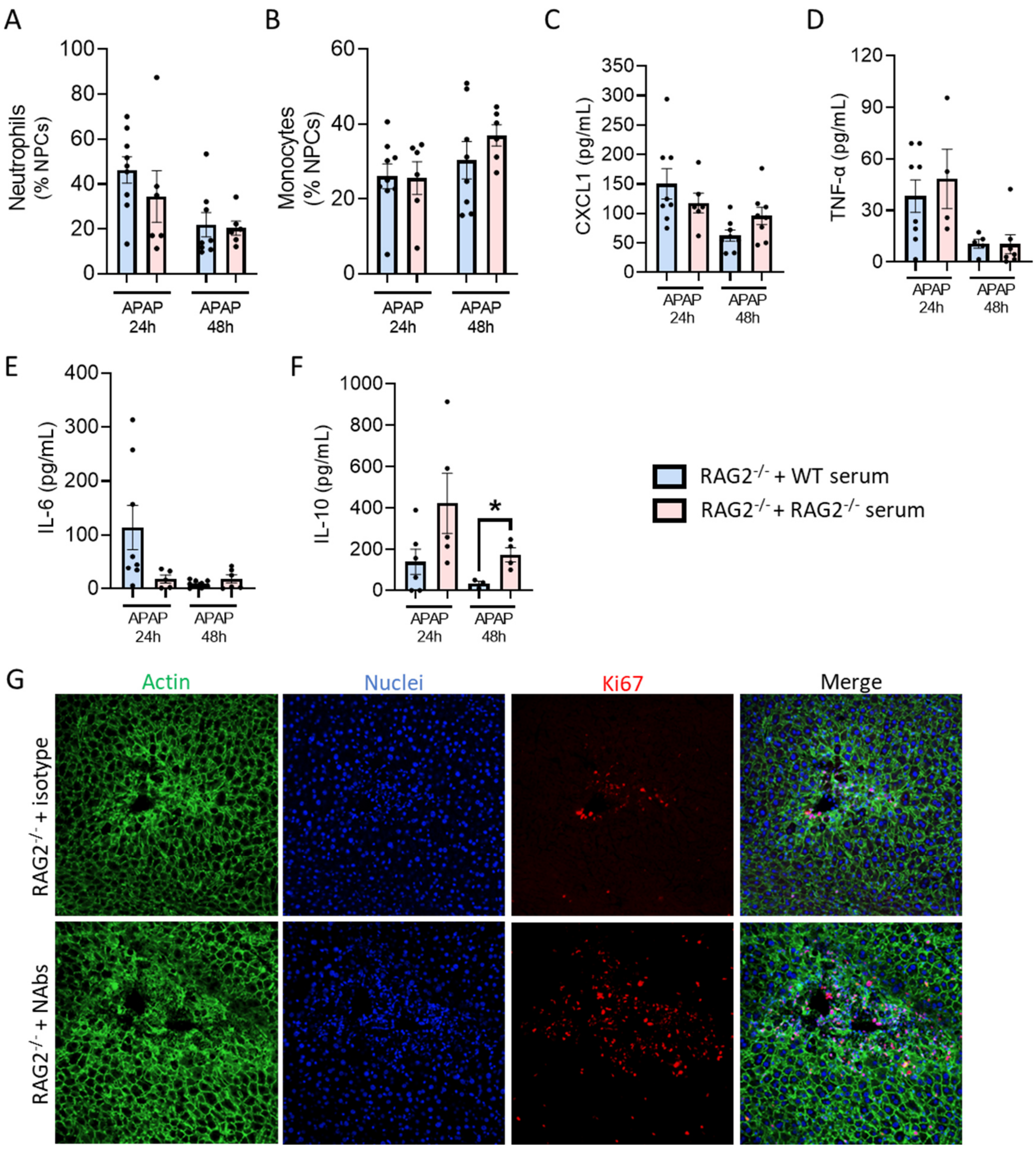


**Supplementary Figure 5:** (A) Flow cytometry of liver non parenchymal cells (NPCs) showing the percentage of neutrophils (Ly6G$^{+}$). (B) Flow cytometry of liver NPCs showing the percentage of monocytes (Ly6G$^{-}$/CCR2$^{+}$). (C-F) Serum levels of (C) CXCL1, (D) TNF-α, (E) IL-6 and (F) IL-10 in RAG2$^{-/-}$ mice after receiving APAP (600 mg/kg) for 24 and 48 hours. (G) Representative images of liver cryosections from RAG2$^{-/-}$ mice 48 hours after receiving APAP showing parenchymal cell proliferation (Ki67$^{+}$, red), F-actin (phalloidin, green) and nuclei (Hoechst, blue). Mice were treated with purified IgM and IgG (100 µg/mouse, i.v.) or isotype. Data is from RAG2$^{-/-}$ mice that received an intravenous injection of WT serum or RAG2$^{-/-}$ serum (A-F) 4 hours after an oral gavage of 600 mg/kg APAP and were sacrificed 24 and 48 hours after the APAP challenge. Each dot in the graph represents a single mouse (A-F). Data are represented as mean ± SEM. *$p < 0.05$.

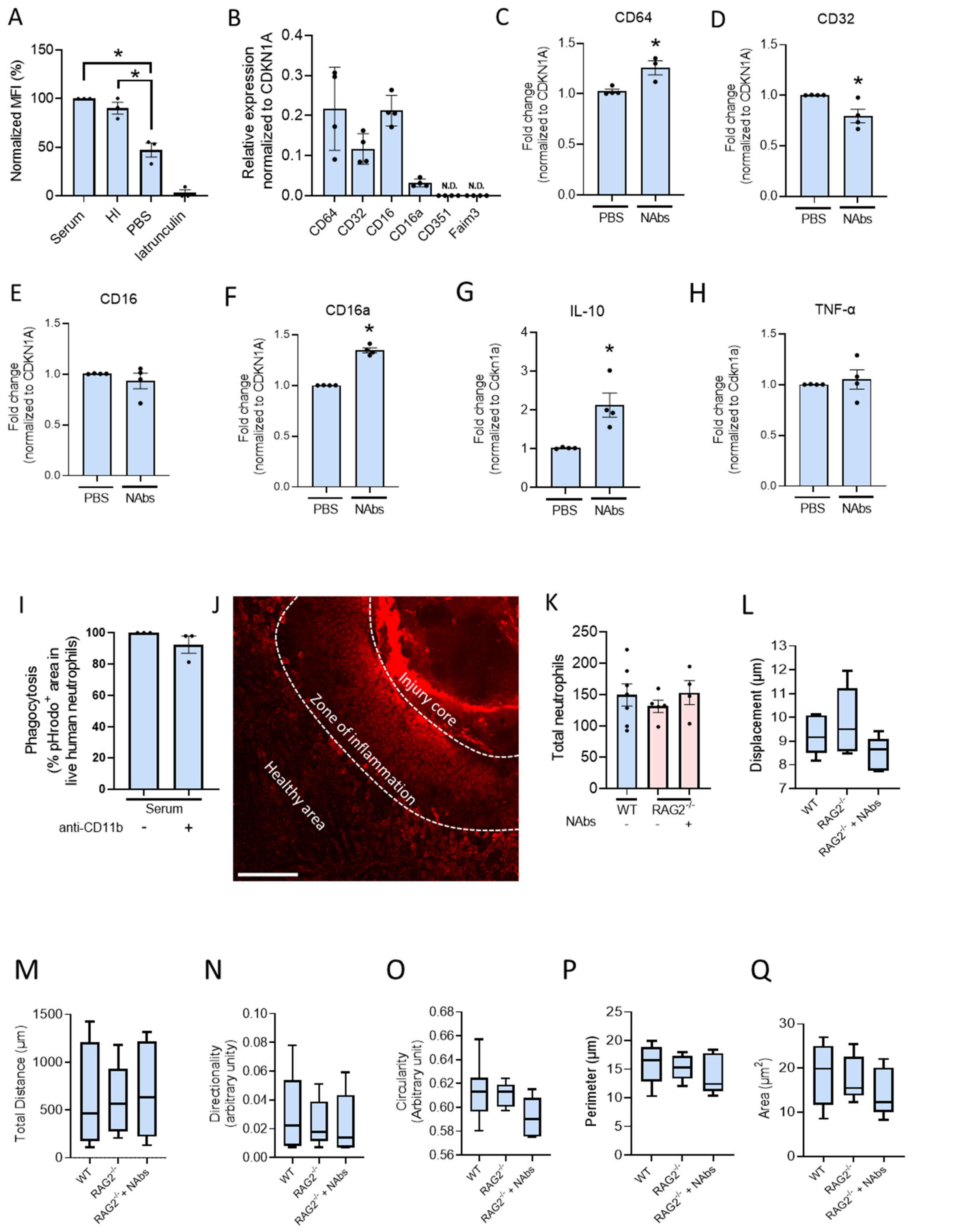


**Supplementary Figure 6:** (A) Mean fluorescence intensity (MFI) of RAW 264.7 cells phagocytosing pHRodo+ debris for 3 hours. Necrotic debris was opsonized with PBS, mouse serum or heat-inactivated serum. Phagocytosis is shown as the normalized percentage compared to the serum group. 10 µM latrunculin was used to block phagocytosis. (B) Gene expression of RAW 264.7 cells in steady state. Data is represented as ΔCT of the target gene over the ΔCT of the housekeeper gene (*Cdnk1a*). CD64 = FcγRI; CD32 = FcγRII; CD16 = FcγRIII; CD16a = FcγRIV; CD351 = Fcα/µ receptor; Faim3 = Fcµ receptor.

(C-H) Gene expression of (C) CD64 (FcγRI), (D) CD32 (FcγRII), (E) CD16 (FcγRIII), (F) CD16a (FcγRIV), (G) IL-10 and (H) TNF-α in RAW 264.7 cells incubated with PBS-opsonized debris or NAbs-opsonized debris (10 µg/ml purified IgM/IgG). Data is represented as $2^{-\Delta\Delta CT}$, relative to the control. *Cdnk1a* was used as a housekeeping gene. (I) Phagocytosis of necrotic debris opsonized with normal serum by primary human neutrophils for 3 hours. CD11b was blocked with 10 µg/ml anti-CD11b. Data is shown as the relative percentage of phagocytosis compared to serum. (J) A representative confocal image of the focal burn injury (after 6 hours) labeled with a droplet 4 µM pHrodo Red succinimidyl ester (red) indicating 3 different zones in the injury. The phagocytosis was imaged in the zone of inflammation. Scale bar = 200 µm. (K) Total number of neutrophils 6 hours after a focal burn injury in the liver of WT, $RAG2^{-/-}$ and $RAG2^{-/-}$ mice injected with 100 µg purified IgG and IgM. Neutrophil numbers were counted manually in one video per mouse. (L-Q) Neutrophil tracking and morphology in WT, $RAG2^{-/-}$ mice and $RAG2^{-/-}$ mice injected with 100 µg purified IgG and IgM. Parameters were quantified with the plugin TrackMate 7 in FIJI ($n \geq 4$ per group). Each dot in the graph represents a single mouse (K-Q). Data are represented as mean ± SEM. *$p < 0.05$.

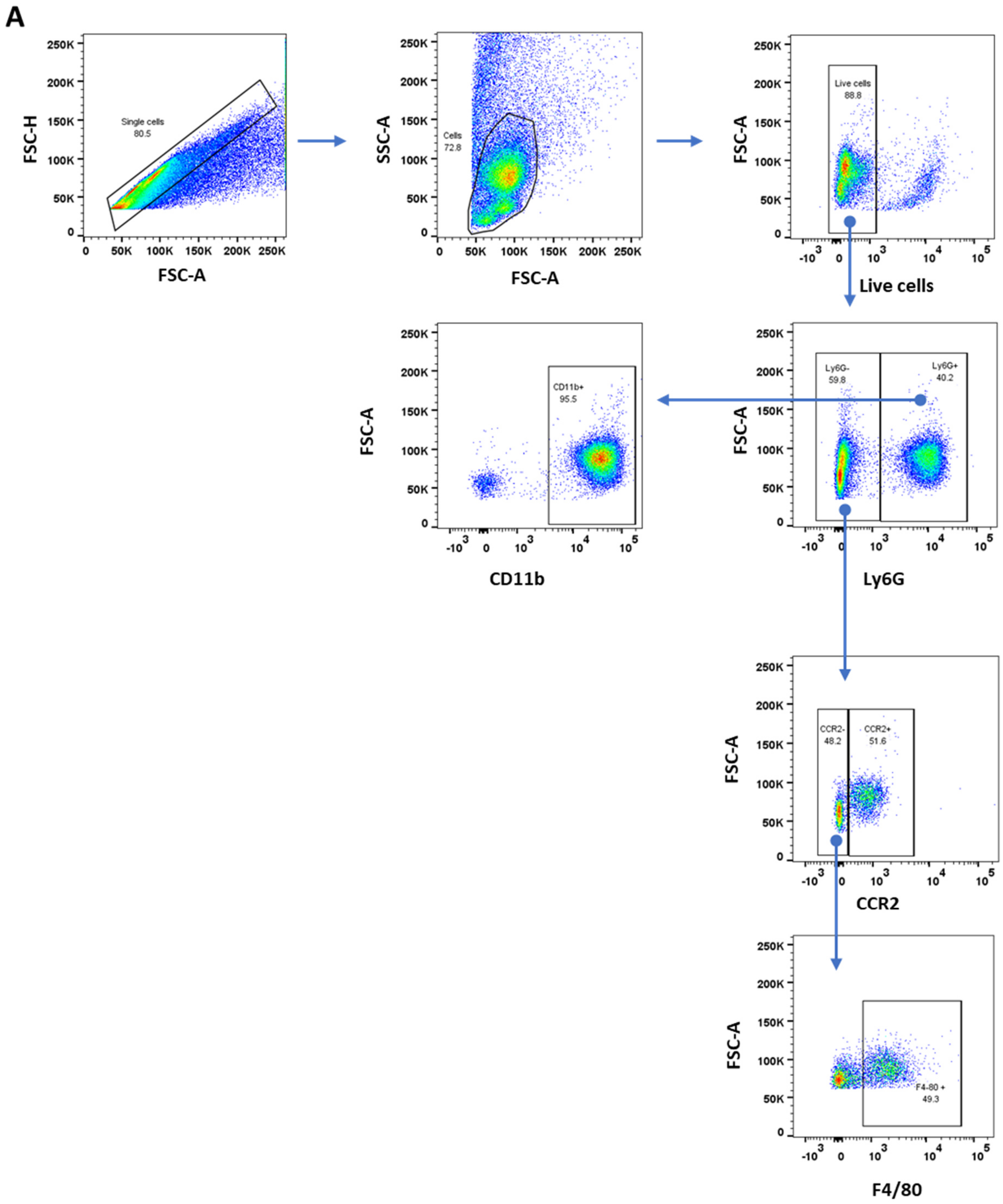


**Supplementary figure 7:** (A) Gating strategy for liver non parenchymal cells.
